# NaP-TRAP: A versatile and accessible workflow to dissect principles of translational regulation and mRNA stability

**DOI:** 10.64898/2026.04.12.718002

**Authors:** Amit Gupta, Anna Z. Struba, Srihari Madhavan, Ethan Strayer, Jean-Denis Beaudoin

**Affiliations:** Department of Genetics and Genome Sciences, UConn Health, Farmington, CT, USA; Binghamton University, State University of New York (SUNY), Binghamton, NY, USA; Department of Genetics, Yale University School of Medicine, New Haven, CT, USA

**Author notes:** Corresponding author, 400 Farmington Avenue, Farmington, CT 06030-6403. These authors contributed equally (Amit Gupta, Anna Z. Struba, Srihari Madhavan).

**Keywords:** Translation, Post-transcriptional regulation, Massively Parallel Reporter Assay, Gene expression

## Abstract

The translation of mRNA into protein is tightly regulated by both cellular *trans*-factors and *cis*-regulatory elements encoded within transcripts. Although transcript fate can be measured by transcript abundance or translation efficiency, separating the contribution of each individual *cis*-element within a single transcript is an ongoing challenge. Current massively parallel reporter assay (MPRAs) approaches enable systematic interrogation of *cis*-regulatory elements that control transcript stability, but translation-focused MPRAs remain technically limited and often inaccessible. Here we present Nascent Peptide Translating Ribosome Affinity Purification (NaP-TRAP), a reporter-based approach that simultaneously measures translation and mRNA abundance. Unlike previous methods, NaP-TRAP captures translation directly through the immunoprecipitation of epitope-tagged nascent peptide chains, providing instantaneous, frame-specific readouts without specialized instrumentation. The method is highly scalable from single reporters to complex libraries, and adaptable across *in vivo* and *in vitro* systems. NaP-TRAP is versatile, allowing assessment of *cis-*regulatory impact of elements distributed throughout the mRNA, from cap-to-tail. This protocol covers experimental design, reporter construction, sample processing, and computational analysis for both low- and high-throughput applications. Bench work can be completed in 4– 5 days, with qPCR-based readouts requiring only basic Excel skills for data processing. Sequencing-based readouts require skills in command-line tools and Python scripting and add an additional 2–3 days. NaP-TRAP thus offers an accessible, robust, and quantitative platform to decode the regulatory logic of mRNA translation and stability in diverse biological contexts.

**Basic Protocol 1:** Design, assembly, and synthesis of NaP-TRAP reporter libraries.

**Support Protocol 1:** Design, assembly, and synthesis of NaP-TRAP individual reporters and spike-ins.

**Basic Protocol 2:** NaP-TRAP delivery by micro-injection in zebrafish embryos.

**Alternate Protocol 1:** NaP-TRAP delivery by transfection in cultured mammalian cells.

**Basic Protocol 3:** NaP-TRAP pulldown and RNA extraction.

**Basic Protocol 4:** Preparation of NaP-TRAP cDNA Sequencing Libraries.

**Alternate Protocol 2:** NaP-TRAP-qPCR module for low-cost validation.

**Basic Protocol 5:** Computational analysis of NaP-TRAP MPRA data.

## INTRODUCTION

The translation of mRNA to proteins is a fundamental process for life, and any misregulation can lead to aberrant protein production (Gingold and Pilpel, 2011), resulting in developmental disorders (Kelleher and Bear, 2008), neurological disorders (Cleary and Ranum, 2013), and cancer (Fabbri et al., 2021). The regulation of translation is primarily managed through the interaction between *trans*-acting factors and *cis*-regulatory elements. *Trans*-acting factors, such as RNA binding proteins (RBPs) (Hentze et al., 2018; Fagre and Gilbert, 2024) and non-coding RNAs (ncRNAs) (Shi et al., 2015), influence the translation machinery from initiation to termination. The recruitment of these *trans*-factors is largely dictated by the presence of *cis*-elements encoded within the transcript itself (Medina-Muñoz et al., 2021). These *cis*-elements can occur in many variations and combinations across the transcriptome, allowing intricate and adaptable control over how different transcripts will be handled in a given cellular context. Often enriched in untranslated regions (UTRs), *cis*-elements can range from short sequences of just a few nucleotides to complex RNA structures spanning several hundred nucleotides (Gupta and Beaudoin, 2021; Byeon et al., 2021). Due to this diversity, their identification and characterization throughout the transcriptome is a significant and evolving challenge.

As a result, many techniques have been developed to study translational regulation at the transcriptome level, including polysome profiling (Chassé et al., 2017; Zuccotti and Modelska, 2016), Translating Ribosome Affinity Purification (TRAP-seq) (Heiman et al., 2008), and ribosome profiling (Ingolia et al., 2009, 2019). These methods have led to groundbreaking discoveries about the translatome and have identified novel translational regulatory patterns across various biological contexts (Fabbri et al., 2021; Reimão-Pinto et al., 2023; Kingston et al., 2022). However, these approaches cannot resolve the effects of individual *cis*-elements, instead capturing the aggregated effect of all regulatory elements on transcript fate. This distinction is important for pathway-level characterization, as endogenous mRNAs frequently contain multiple *cis*-regulatory elements with context-dependent and potentially competing functions. An approach to isolate these *cis*-elements while assessing their impact on translation is needed.

The standard for identifying and characterizing mRNA *cis*-regulatory elements has been the use of reporter genes (Akirtava and McManus, 2021; Dao et al., 2025; Mikl et al., 2022; Rabani et al., 2017). Recent advancements in DNA oligo synthesis have revolutionized reporter assay approaches by significantly enhancing their throughput, leading to the development of massively parallel reporter assays (MPRAs). These MPRAs utilize complex reporter libraries to evaluate the regulatory activity of thousands of mRNA sequences on specific mRNA functions, such as stability (Siegel et al., 2022) and translation (Reimão-Pinto et al., 2023). By varying only a segment of the reporter mRNA while keeping all other components constant, MPRAs focus on analyzing the elements within these variable regions, thereby eliminating the confounding effects of other elements present in endogenous transcripts. In the realm of translation, MPRA methodologies such as growth selection (Cuperus et al., 2017), FACS (Gilliot and Gorochowski, 2023), polysome profiling (Reimão-Pinto et al., 2023; Zhao et al., 2014), and Direct Analysis of Ribosome Targeting (DART) (Niederer et al., 2022) (Table 1) have offered a comprehensive analysis of known elements and uncovered new sequence motifs that drive translation in mRNA 5’- and 3’-UTRs. Although these tools have yielded significant insights into the regulation of translation, their widespread use has been constrained by technical limitations and the requirement for specialized equipment (see Table 1).

**Table 1.** Characteristics of current techniques for evaluating the regulatory impact of mRNA *cis-*regulatory elements on translation.

| Method | System | Specialized Equipment required | Frame-specific | Translation Read-out | Source of signal |
| --- | --- | --- | --- | --- | --- |
| Growth-selection Assays | Cell lines | None | Yes | Steady-State | Accumulated proteins (transcription & translation) |
| FACS | Cell lines | Fluorescent cell sorter | Yes | Steady-State | Accumulated proteins (transcription & translation) |
| DART-Seq | Cell lysate | Ultra-centrifuge, Sucrose Fractionator | Yes (reporter can only contain 1 ORF) | Equilibrium | Single bound ribosome at AUG |
| Polysome profiling | Model organisms & cell lines (High input) | Ultra-centrifuge, Sucrose Fractionator | No | Instantaneous (snapshot) | Bound ribosomes |
| NaP-TRAP | Model organisms & cell lines (Low input) | None | Yes | Instantaneous (snapshot) | Translating ribosomes (Tag-containing ORF) |

To address the limitations of current MPRA methods for studying translational regulation, we aimed to develop a method that is both simple and adaptable, allowing for broad application across various systems without requiring specialized equipment. We recognized that immunocapture is a rapid and widely used technique in various biological fields and therefore made it the foundation of this new MPRA, NaP-TRAP (Nascent Peptide Translating Ribosome Affinity Purification) (Fig. 1A). NaP-TRAP relies on the insertion of an N-terminal epitope tag into the reporter open reading frame. This approach effectively tagged the nascent peptide of actively translating ribosomes, enabling the immunoprecipitation of mRNA reporters based on the number of ribosomes engaged in translation (Fig. 1B). This design ensures specific enrichment of actively translating ribosomes in a frame-specific manner. Moreover, NaP-TRAP offers a real-time snapshot of translation, making it ideal for examining dynamic processes where protein production changes rapidly.

**Figure 1.**
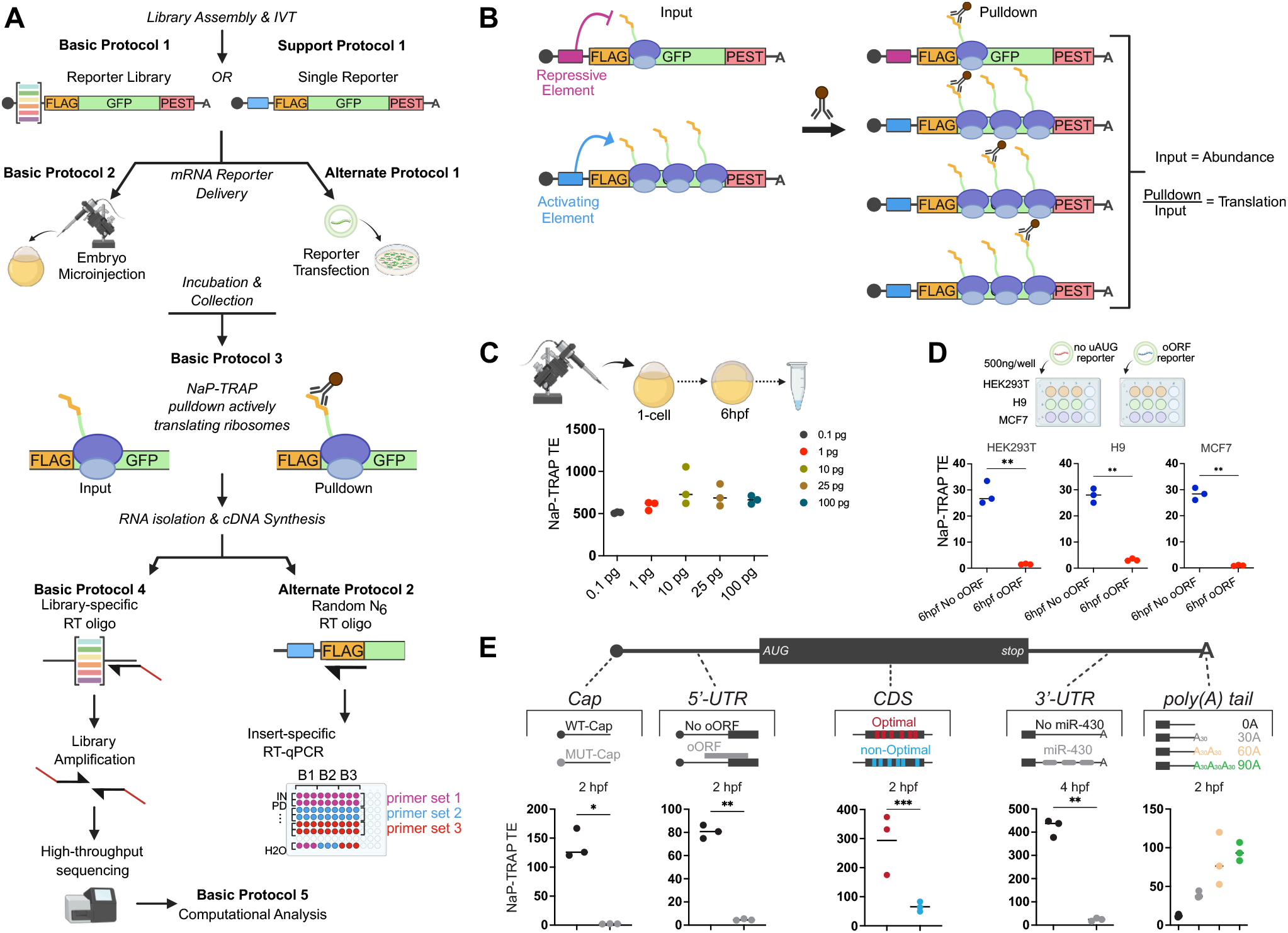
Overview of the NaP-TRAP workflow and examples of use cases. (**A**) NaP-TRAP can be performed using either complex reporter libraries or individual reporters, delivered *in vivo* by zebrafish embryo microinjection or *in vitro* by mammalian cell transfection, followed by pulldown, RNA purification, and either sequencing- or qPCR-based readout. (**B**) Principle of NaP-TRAP: FLAG-tagged nascent peptides on actively translating ribosomes are immunocaptured to enrich ribosome-associated reporter mRNAs (Pulldown). Reporter abundance is measured in the Input, and translation output is quantified as Pulldown/Input (NaP-TRAP TE). (**C**) Robust delivery and readout across a 100-fold range of injected reporter amounts in zebrafish embryos, with consistent NaP-TRAP TE across all doses. (**D**) Comparable reporter trends across mammalian cell types illustrated by differential translation of control versus oORF-containing reporters in HEK293T, H9, and MCF7 cells. (**E**) NaP-TRAP supports interrogation of diverse regulatory features across the reporter mRNA, including mRNA cap types, 5′-UTR elements (e.g., uORFs/oORFs), coding sequence codon optimality, 3′-UTR elements (e.g., miR-430 sites), and poly(A) tail length in zebrafish embryos. Cartoon diagrams were created individually in BioRender (Smith, J. (2025). BioRender.com/c248457).

NaP-TRAP can be assessed through RT-qPCR or Illumina sequencing to determine the abundance of reporters in both input and pulldown fractions, contingent on the number of reporters involved (Fig. 1A). We have verified that the pulldown-to-input ratio reliably indicates the translational status of each reporter at the moment of capture (Strayer et al., 2024). For example, we were able to resolve the temporal repression mediated by miR-430 during the maternal-to-zygotic transition (MZT) in zebrafish (Strayer et al., 2024). We also confirmed that NaP-TRAP-derived translation values are not biased by mRNA abundance, as normalization to input levels enables fair comparisons across low- and high-expression reporters (Fig. 1C). Furthermore, we demonstrated that NaP-TRAP is highly scalable, delivering accurate translation values whether using a single reporter or more than 10,000 reporters (Strayer et al., 2024). Finally, we successfully applied NaP-TRAP in a range of *in vitro* and *in vivo* systems (Fig. 1D) (Strayer et al., 2024), highlighting its adaptability for studying translation across biological contexts.

NaP-TRAP does not require any specialized equipment but instead relies on immunocapture by magnetic beads. This pulldown strategy is widely used in research labs, making NaP-TRAP appealing to scientists interested in translational regulation. The versatility of NaP-TRAP in adapting to various cell lines and *in vivo* models enhances its applicability across numerous systems and research fields (Fig. 1C-D). We showed that NaP-TRAP can measure the regulatory activity found anywhere in an mRNA, from cap-to-tail, making it suitable for investigation of a wide diversity of regulatory elements (Fig. 1E). For example, NaP-TRAP can identify regulatory elements within the coding sequence, such as codon optimality (Bazzini et al., 2016), as well as within UTRs such as uORFs and miRNA binding sites (Fig. 1E). As further evidence, the Bartel lab used NaP-TRAP to uncover 3’-UTR *cis*-regulatory elements of translation in frog oocytes (Xiang et al., 2024). NaP-TRAP can also be paired with RBP perturbations to dissect mechanisms. For instance, in a collaboration with the Bushell lab, we combined NaP-TRAP and distinct eIF4A1 inhibitors to reveal feature-specific deployment of eIF4A1 functions and regulon selection (Schmidt et al., 2025). Finally, NaP-TRAP not only measures the impact of *cis*-elements on translation, but also allows for the simultaneous assessment of mRNA stability through the analysis of input values, all within a single experiment.

Because NaP-TRAP can resolve reading frames, it can be applied to a broad range of frame-specific applications. For instance, we demonstrated that it is possible to introduce three distinct tags, one in each of the three reading frames, within a reporter. This allows us to measure the translation occurring in each frame at a specific time within the same experiment (Strayer et al., 2024). By employing similar manipulations of various tags, NaP-TRAP can be utilized to investigate IRES activity in bicistronic reporters, frameshifting elements, or translation readthrough. This expands its scope and establishes it as a powerful tool for gaining a deeper understanding of translation regulation.

We have initially focused on a reporter-based approach, but the underlying principle of the method suggests it could also be applied to measure translation of endogenously tagged mRNAs. With the continued advancement of genome editing technologies, it is increasingly feasible to insert epitope tags at precise genomic locations. For instance, tagging the N-terminus of an endogenous gene could make it detectable by NaP-TRAP. We are currently evaluating this strategy *in vivo* in zebrafish using the GEARs toolkit (Boswell et al., 2025).

The first generation of MPRAs investigating translation relied on growth selection and fluorescence-activated cell sorting (FACS) to detect changes in reporter abundance. These assays measured long-term accumulation of proteins or enrichment in cell populations with different levels of fluorescent proteins as a proxy for translation control. A key limitation of these pioneering methods is that they reflect steady-state protein levels, rather than capturing translation dynamics in realtime. Consequently, they are not well-equipped to investigate rapid or temporal translational regulation. Additionally, they fail to differentiate between translation effects and variations in mRNA levels or stability.

To address these limitations, more recent MPRA techniques like polysome profiling (Chassé et al., 2017; Zuccotti and Modelska, 2016) and DART (Niederer et al., 2022) have been introduced. These techniques isolate mRNAs based on ribosome association, offering a snapshot of translation at a specific time point (Table 1). When we benchmarked NaP-TRAP against polysome profiling, we observed a strong overall correlation (R^2^ = 0.85) between mean ribosome load and NaP-TRAP-derived translation values (Strayer et al., 2024). However, we also noted that polysome profiling tends to overestimate translation of reporters containing upstream AUGs that form upstream ORFs (uORFs) or overlapping ORFs (oORFs) (Johnstone et al., 2016; Calvo et al., 2009; Meijer and Thomas, 2002). This overestimation likely arises from the fact that techniques such as polysome profiling and TRAP-seq measure total ribosome occupancy, encompassing inactive ribosomes or those translating regions out of frame, rather than focusing exclusively on the main coding sequence. In contrast, NaP-TRAP captures translation specifically within a defined ORF, providing a frame-specific and real-time measure of protein synthesis. The ability to insert distinct tags in separate frames further enables NaP-TRAP to measure translation in multiple reading frames within a single experiment, which is an approach not possible with other methods.

NaP-TRAP provides several key advantages over more complex techniques like FACS (Gilliot and Gorochowski, 2023) and polysome profiling. First, it is well suited for low-input applications, yielding reproducible translation values from as little as 0.1 picogram of injected mRNA (Fig. 1C). Second, NaP-TRAP’s straightforward input normalization generates a continuous and easily interpretable translation metric, in contrast to normalization schemes required by gradient fractionation (Chassé et al., 2017; Zuccotti and Modelska, 2016) or FACS (Gilliot and Gorochowski, 2023). Third, because of this normalization, NaP-TRAP translation values remain consistent across a 100-fold range of reporter mRNA levels (Fig. 1C), highlighting its robustness. Fourth, NaP-TRAP can be used in systems with low global translation or where monosomes dominate, such as early embryos or neurons. Fifth, unlike methods that require ultracentrifuges or flow cytometers, NaP-TRAP does not necessitate specialized equipment. DART (Niederer et al., 2022) (Table 1) is a MPRA that estimates translation by incubating reporters in translation-incompetent lysates, followed by ultracentrifugation to separate 80S ribosome-bound mRNAs from unbound transcripts. Translation is quantified by determining a ribosome recruitment score, which is calculated from the ratio of bound reporter levels to those of the input. While DART’s simplicity enhances both sensitivity and reproducibility, it differs from NaP-TRAP in several significant ways. DART necessitates ultracentrifugation, is limited to *ex vivo* analyses, and is ineffective in assessing the translation of reporters with upstream AUGs or multiple reading frames.

To date, all NaP-TRAP experiments have been conducted using *in vitro* transcribed mRNA reporters. This approach has several advantages, such as eliminating concerns related to transcriptional artifacts like unintended promoter activity, alternative transcription start sites (TSSs), or cryptic splice site usage. However, it also presents important limitations. Because the RNA is synthesized *in vitro*, it lacks endogenous RNA modifications that are deposited co-transcriptionally, such as N6-methyladenosine (m6A). Although canonical nucleotides can be substituted with modified analogs (*e*.*g*., N1-methylpseudouridine to reduce immunogenicity in mRNA vaccines), introducing specific site-directed modifications remains technically challenging. Furthermore, the function of certain *cis*-regulatory elements may depend on nuclear *trans*-factors or nuclear processes like splicing and mRNA export. These elements are likely to be inaccurately assessed when mRNAs are delivered directly into the cytoplasm, bypassing nuclear processing. To address these limitations, we are currently developing a version of NaP-TRAP compatible with DNA-based reporters. This will more accurately recapitulate endogenous gene expression, enabling regulatory elements to engage with nuclear factors and processes. Simultaneously, we are developing a sequencing library construction approach aimed at minimizing unintended pre-mRNA processing events, such as cryptic splicing or alternative transcription start sites, which could otherwise complicate interpretation.

Another early concern was whether accumulation of full-length tagged proteins over time could interfere with NaP-TRAP’s ability to measure ongoing translation. Since these proteins share the same tag as the nascent peptides on translating ribosomes, they might compete for binding to the antibody-coated beads, potentially narrowing the window in which NaP-TRAP can accurately capture a snapshot of translation. To mitigate this, we incorporated a short PEST domain at the C-terminus of the tagged proteins. This degron significantly reduced protein half-life in both zebrafish embryos and various mammalian cell lines, thereby minimizing interference from mature protein during immunoprecipitation. However, in systems where the PEST domain is less active or functionally impaired, this issue may re-emerge and will require system-specific validation or alternative degradation strategies.

This article describes the complete NaP-TRAP workflow, organized into five Basic Protocols. **Basic Protocol 1** describes the design and assembly of reporter libraries from oligo pools, and **Support Protocol 1** outlines a strategy to synthesize individual reporters and spike-ins controls. **Basic Protocol 2** and **Alternate Protocol 1** describe the delivery of reporter constructs into zebrafish embryos or mammalian cells, respectively. Following delivery, **Basic Protocol 3** outlines sample collection and NaP-TRAP pulldown. Pulldown and input samples can then be processed for either high throughput sequencing (**Basic Protocol 4)** or for qPCR-based analysis (**Alternate Protocol 2)**. Finally, **Basic Protocol 5** describes processing and analysis of NaP-TRAP-sequencing datasets, including a detailed overview of the associated computational pipeline.

### CAUTIONS

All chemical reactions with hazardous chemicals (*e*.*g*., cycloheximide, phenol, chloroform, TRI reagent, EtBr, TRIzol) must be performed under a chemical fume hood with proper eye protection, gloves, and lab coat. All liquid waste containing phenol and/or chloroform should be disposed of as per EHS regulatory guidelines. Ethanol and isopropanol are highly flammable and must be stored and handled with caution.

Fish lines are maintained following the International Association for Assessment and Accreditation of Laboratory Animal Care research guidelines protocol approved by University Connecticut Health Institutional Animal Care and Use Committee (IACUC). Fish are maintained at 28°C on a 14-h light/10-h dark cycle. Wild-type zebrafish embryos are obtained through natural mating of TU, AB, TUAB, TL or NHGRI strains of mixed ages (5-18 months). All embryos are kept in system water with methylene blue. Dechorionated embryos are kept on agar plates made with methylene blue system water. Minimize the time that embryos spend outside of 28°C conditions; developmental timepoints can fluctuate with changes in temperature, and must be tracked closely.

### STRATEGIC PLANNING

The full NaP-TRAP workflow (Fig. 1A) can be completed by an experienced technician within one week depending on the readout strategy, but requires planning to ensure efficiency and success.

Reporter assembly and IVT can be completed within 1 day for oligo pools or non-cloned constructs, but an extra 2 days are required if the additional bacterial cloning steps are performed (see **Basic Protocol 1** and **Support Protocol 1**). Cloning individual reporter constructs is convenient for small-scale experiments and for generating spike-in controls, but requires sequence validation by Sanger sequencing. In parallel with PCR and IVT to generate NaP-TRAP reporter mRNAs, you can begin preparing embryos or cultured cells.

Zebrafish crosses require overnight separation of males and females. Because fertility can vary, plan to inject at least 30% more embryos than you intend to collect to ensure sufficient numbers of fertilized embryos at the desired stage (**Basic Protocol 2**). Seeding mammalian cells for construct delivery may take several days to achieve optimal density and cell health before transfection (**Alternate Protocol 1**).

Post-collection of samples, NaP-TRAP pulldown can be completed quickly in ∼3 hours (**Basic Protocol 3**), but plan to complete subsequent RNA isolation and cDNA synthesis on the same day to minimize degradation (**Basic Protocol 4 or Alternate Protocol 2**). For NaP-TRAP-qPCR (**Alternate Protocol 2)**, reverse transcription can be performed using random hexamers, whereas NaP-TRAP-sequencing (**Basic Protocol 4**) requires library-specific primers with barcoded replicates (see Fig. 2F).

**Figure 2.**
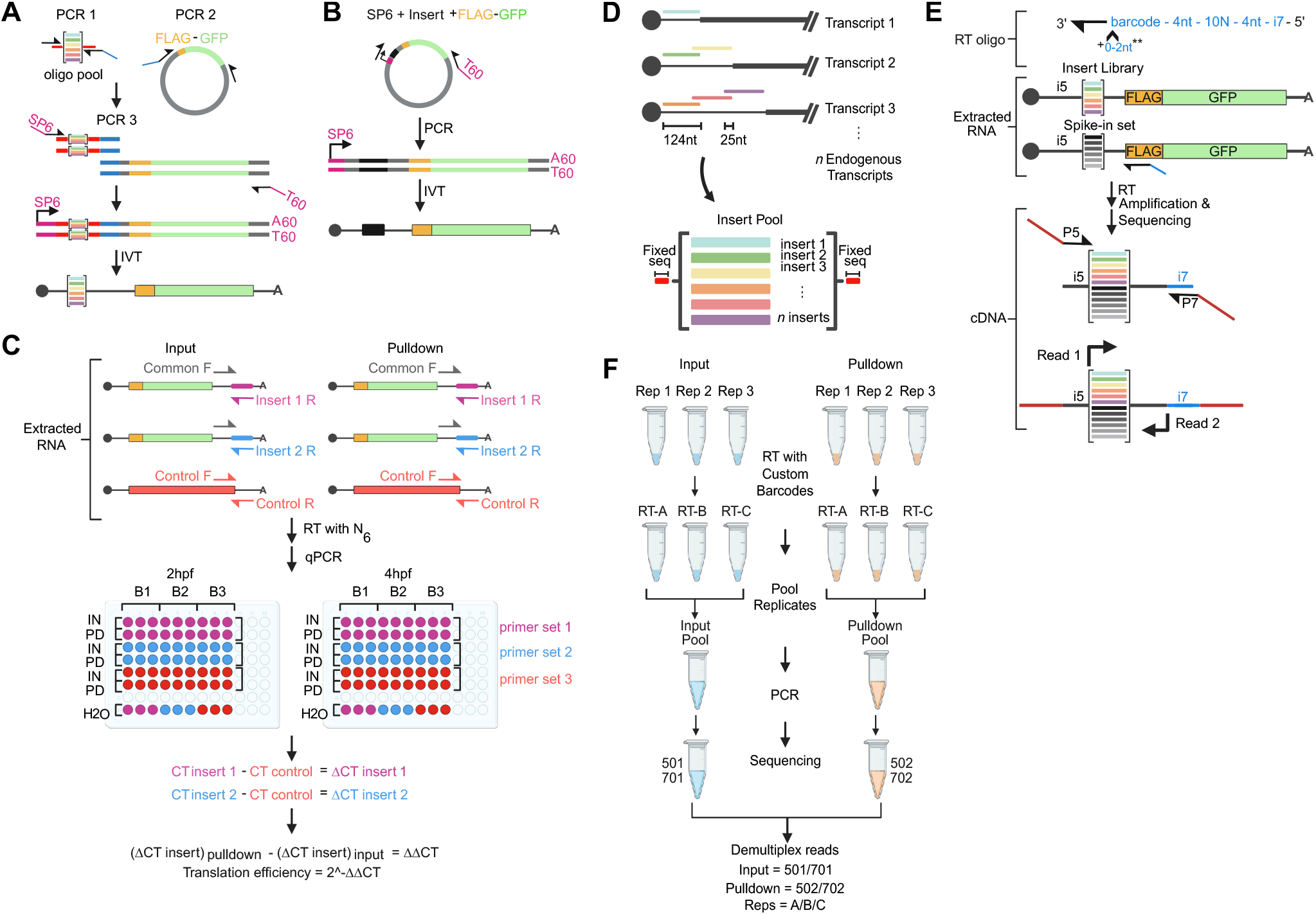
NaP-TRAP reporter assembly and sequencing library preparation. (**A**) Complex NaP-TRAP reporter libraries for MPRA applications are generated by PCR-based assembly, in which pooled inserts are fused to the shared FLAG–GFP reporter backbone and flanking sequences required for *in vitro* transcription. (**B**) Single reporters can be cloned individually and linear *in vitro* transcription templates produced by PCR using primers that add the appropriate promoter and hard-encoded poly(A) tail. (**C**) For qPCR readout, reporter abundance in matched Input and Pulldown fractions is quantified using insert-specific primer sets and normalized to a co-delivered control reporter (e.g., dsRed). (**D**) Example design of a tiled 5′-UTR library, in which endogenous sequences are sampled using 124-nt windows with 25-nt steps. (**E**) For sequencing readout, NaP-TRAP RNA is reverse transcribed using a custom barcoded RT primer that incorporates a sample barcode, a 10N unique molecular identifier (UMI), and an i7 index sequence. Subsequent PCR-amplification with indexed Illumina primers yields a dual-indexed sequencing library compatible with Illumina platforms. (**F**) Reverse-transcribed samples with distinct custom barcoded RT primers can be pooled prior to the final PCR amplification with indexed Illumina primers. This pooling step promotes uniform PCR amplification across replicates. After sequencing, reads from each replicate are recovered during demultiplexing using the RT primer barcode.

Once samples are reverse transcribed, the remaining workflow takes 1-4 days, depending on the readout. For qPCR–readout, reporter-specific primers should be designed in advance, along with a control primer set for a co-delivered normalization reporter (**Alternate Protocol 2**). qPCR can typically be completed and analyzed the same day. For sequencing-readout, cDNA is amplified using primers that add Illumina-compatible indices (**Basic Protocol 4**). The resulting libraries should first be evaluated by small-scale sequencing (e.g., MiSeq) to confirm complexity and diversity before scaling to a higher-throughput platform such as NovaSeq if needed, adding an additional 2-3 days. Post deeper sequencing, computational analysis of NaP-TRAP-sequencing datasets generally requires 6-7 hrs, depending on available resources (**Basic Protocol 5**).

#### Basic Protocol 1: DESIGN, ASSEMBLY, AND SYNTHESIS OF NaP-TRAP REPORTER LIBRARIES

This protocol details how to design, assemble, and synthesize NaP-TRAP reporter libraries starting from an oligo pool of 5’-UTR regions as an example. Library construction uses a three-step PCR workflow (Fig. 2A): PCR1 amplifies the oligo pool and appends the required flanking sequences; PCR2 amplifies the constant NaP-TRAP reporter core, including the FLAG encoding coding sequence; and PCR3 fuses the PCR1 and PCR2 products to generate a full-length DNA template bearing an SP6 promoter at the 5′ end and a 3′ hard-encoded poly(A) tract compatible with *in vitro* transcription. *In vitro* transcription then produces a capped NaP-TRAP mRNA library in which every reporter shares the same FLAG-GFP coding sequence and 3′-UTR, while differing only in the variable 5′-UTR region (Fig. 2A). When performed correctly, the procedure yields a high-complexity mRNA reporter library suitable for downstream NaP-TRAP translation measurements.

#### Materials

Plasmid templates (see **Supporting Information**: Table S1 for all sequences)

*All the plasmid constructs used in this study were cloned in pCS2+ vector which was generously gifted by Antonio Giraldez lab*.

Nuclease-free water (Sigma-Aldrich, Cat. No. W4502-1L)

1 M TRIS (Thermo Fisher Scientific, Cat. No. 15567027)

KAPA HiFi HotStart Polymerase Readymix (Roche, Cat. No. 07958927001)

Oligos (see **Supporting Information**: Table S1) ordered from Sigma or IDT with standard desalting

Oligo pools from GenScript and Twist Bioscience

Thermal cycler (BIO-RAD, Model No. S1000)

DNA Clean & Concentrator (Zymo, Cat. No. D4006)

6X Gel loading dye (New England Biolabs, Cat. No. B7025)

EtBr (BIO-RAD Cat. No. 1610433)

1 kb Plus DNA Ladder (New England Biolabs, Cat. No. N3200)

TAE buffer (see **REAGENTS AND SOLUTIONS**)

Agarose gel 1% (wt/vol, see **REAGENTS AND SOLUTIONS**)

mMessage mMachine SP6 RNA Polymerase kit (Thermo Fisher Scientific, Cat. No. AM1340)

DNase I (New England Biolabs, Cat. No. M0303S)

RNA Clean & Concentrator-5 (Zymo, Cat. No. R1013)

Filter-tips 20 µL, 200 µL and 1,000 µL (RAININ, Cat. Nos. 30389225, 30389240, 30389212)

Micro Centrifuge tubes 1.5 mL and 2 mL (USA Scientific, Cat. Nos. 1615-5510 and 1420-2700)

Low-binding reaction tubes 1.5 mL (Eppendorf, Cat. No. 022431021)

8 PCR tube stripes 0.2 mL (USA Scientific, Cat. No. 1402-4700)

NanoDrop (Fisher Scientific, Cat. No. 13-400-526)

Gel electrophoresis system (*e*.*g*., Thermofisher Scientific, A2-BP)

Gel Imager (BIO-RAD, Model No. ChemidocTM MP Imaging System)

Blue light gel transilluminator (CLARE CHEMICAL RESEARCH DR46B)

Zymoclean Gel DNA Recovery Kit (Zymo, Cat. No. D4007)

Heat block (*e*.*g*., Thermofisher Scientific, Cat. No. 88870003)

Table-top Centrifuge Refrigerated (Eppendorf, Model No. 5427R)

#### Prepare oligo insert library for assembly

1. Design and order the oligo pool for reporter inserts with flanking fixed sequences following considerations mentioned in the ***Reporter design: Basics*** and ***Reporter design: Illumina sequencing readout*** sections (Fig. 2A).
2. Resuspend the oligo pool to 10 ng/µL in Nuclease-Free Water (*e*.*g*., To a lyophilized library of 100 ng, add 10 µL of Nuclease-Free Water).
3. Vortex thoroughly, and store in aliquots of 3 µL at -20°C.
4. Prepare the following PCR mastermix: 12.5 µL 2X KAPA HiFi HotStart ReadyMix 0.75 µL SP6_fwd (10 µM) 0.75 µL GFP-aug_rev (10 µM) 1 µL Reporter Oligo Pool at 10 ng/µL 10 µL Nuclease-Free Water (NFW) *We recommend the KAPA HiFi HotStart enzyme for all assembly PCRs for its high-fidelity and low amplification bias across GC content, providing better library uniformity*.

5. Mix the reaction and run the following program in a thermocycler:

3 min 95°C

10-14 cycles: 20 s at 98°C

15 s at 60°C

15 s at 72°C

*It is essential to minimize the number of cycles to ensure that each reporter has even representation. Oligo pool cycling should be adjusted based on oligo length, as recommended by the oligo manufacturer. Twist Bioscience recommends 10-12 cycles for 100-150 nt oligo, and 12-14 cycles for 151-300 nt oligos*.

6. Concentrate the reaction using the DNA Clean & Concentrator kit. Elute in 20 µL of Nuclease-Free Water.
7. Add 4 µL of 6X loading dye, and gel-extract the purified reaction using a 2% agarose gel and a Gel DNA Recovery kit. Elute in 12 µL of water and read concentration by NanoDrop. Store at -20°C until needed in future steps.

#### Prepare reporter coding sequence for assembly

8. Amplify the 3XFLAG-GFP-PEST sequence from the plasmid DNA template using the following mastermix: 12.5 µL 2X KAPA HiFi HotStart ReadyMix 0.75 µL 3xFLAG_GFP_fwd (10 uM) 0.75 µL sv40_rev (10uM) 1 µL pCS2+GFP-PEST (diluted to 1ng/uL) 10 µL Nuclease-Free Water

9. Mix, then run the following thermocycler program: 3 min at 95°C 25 cycles: 20 s at 98°C 15 s at 60°C s at 72°C

10. Concentrate the reaction using the DNA Clean & Concentrator kit . Elute in 20 µL of water.
11. Add 4 µL of 6X loading dye, and gel-extract the purified reaction using a 1% agarose gel and a Gel DNA Recovery Kit . Elute in 15 µL of Nuclease-Free Water and measure concentration using NanoDrop. Store at -20°C until needed in future steps.

#### Assemble amplicons

12. Make the overlap extension mastermix with the following reaction (keep on ice until ready to cycle): 25 µL 2X KAPA HiFi HotStart ReadyMix 1 µL Reporter Oligo Pool amplicon (diluted for 1:1 molar ratio with 3XFLAG-GFP-PEST) 1 µL 3XFLAG-GFP-PEST amplicon (diluted for 1:1 molar ratio with Oligo Pool amplicon) 20µL Nuclease-Free Water

13. Prepare the primer mastermix by adding 2 µL of each 100 µM primer to a new tube (do not add it to the PCR reaction yet, it should be added after the extension phase). *For the NaP-TRAP 5’UTR library, we used SP6_fwd, and sv40_60A_rev. The reverse primer adds a 60-nt poly(T) sequence into the amplicon, resulting in a 60A poly(A) tail after in vitro transcription. See* ***Reporter Design: Basics*** *for more details*.

14. Complete the following cycling conditions: 3 min at 95°C 10 cycles: 20 s at 98°C 15 s at 71°C 1.5 min (15-60 s/kb) at 72°C Pause: add 2 µL of each 100 µM primer (premixed, 4 µL total volume) 20 cycles: 20 s at 98°C 15 s at 60°C 1.5 min (15-60 s/kb) at 72°C Final extension: 5 min at 72°C Hold: 4°C

15. Concentrate the reaction using the DNA Clean & Concentrator kit and gel extract the correct band size using 1% gel as mentioned above. Elute in 20 µL TV. *For our design, the correct band size is ∼1000bp*.

16. Utilize a NanoDrop to quantify the concentration of the extracted DNA, ensuring it is concentrated and cleaned enough for *in vitro* transcription with the mMessage mMachine kit. *For the following IVT step, 0*.*2 µg is necessary per sample; we typically see 500-1000 ng/µL at this step*.

#### *In vitro* transcription

17. Make mRNA from template reporter DNA using mMESSAGE mMACHINE SP6 Transcription Kit followed by DNase treatment: 5 µL of 2x NTP/CAP 1 µL of 10X Buffer 3.5 µL of DNA template (0.1-0.5 µg) 0.5 µL of SP6 mMessage mMachine Enzyme mix

18. Incubate for 2 hrs at 37°C.
19. Treat with 1 µL of DNase, and incubate for 15 min at 37°C.
20. Concentrate the reaction using an RNA Clean & Concentrator, elute in 20 µL Nuclease-Free Water and measure concentration by NanoDrop.
21. Aliquot into strip tubes and store at -80°C. *For zebrafish micro-injection, we use aliquots of 2 µL at 100 ng/µL TV. On injection day, this can be diluted with 8 µL of water, to generate 20 ng/µL RNA working solution and injected at 20 pg (1 nL) per embryo. For mammalian cell line transfection, we use aliquots of 2 µL at 1000 ng/µL, to directly use the ug of RNA necessary per well*.

22. Thaw one tube to check the quality of the RNA product.
  a. Heat at 65°C for 3 min.
  b. Cool on ice for 2 min before mixing with loading dye and running on a 2% gel.

*There should be a single, clean band with minimal background*.

23. Continue to **Support Protocol 1** for individual reporter assembly, or to **Basic Protocol 2** and **Alternative Protocol 1** to continue to NaP-TRAP delivery.

#### Support Protocol 1: DESIGN, ASSEMBLY, AND SYNTHESIS OF NaP-TRAP INDIVIDUAL REPORTERS AND SPIKE-INS

This support protocol describes how to design, amplify, and synthesize individual NaP-TRAP reporter mRNAs and spike-in controls for targeted validation and normalization (Fig. 2B). Individual reporters and spike-ins are first designed and cloned into plasmids containing the FLAG-GFP coding sequence. They are then amplified in a single PCR using primers that append a SP6 promoter at the 5′ end and a 60-nt poly(A) tract at the 3′ end, generating an *in vitro* transcription-ready DNA template. This PCR product is subsequently used for *in vitro* transcription to produce individual 5′ capped NaP-TRAP reporter or spike-in mRNAs. When performed correctly, the procedure yields NaP-TRAP reporters mRNAs suitable for the NaP-TRAP-qPCR module and spike-in reporters for normalization in the NaP-TRAP-sequencing workflow. Spike-ins need to be synthesized separately from the complex NaP-TRAP libraries because they are added during RNA extraction of the input and pulldown samples to control for variability introduced during the pulldown and downstream processing.

1. Design reporters and/or spike-ins following guide lines found in the **Reporter design: Basics, Reporter design: qPCR readout**, and **Pulldown and spike-ins** sections below.
2. Clone each NaP-TRAP reporter or spike-in into a plasmid following standard bacterial cloning techniques.
3. Amplify the reporter construct from the plasmid template using the same primers and protocol as described in **Basic Protocol 1**. Similarly, gel extract the product and perform IVT as previously described.
4. Continue to **Basic Protocol 2** or **Alternative Protocol 1**.

#### Basic Protocol 2: NaP-TRAP DELIVERY BY MICRO-INJECTION IN ZEBRAFISH EMBRYOS

This protocol describes how to deliver *in vitro*–transcribed NaP-TRAP reporter mRNAs (individual reporters or complex libraries) into zebrafish embryos by micro-injection. Adult fish are set up for crosses, embryos are collected and dechorionated with pronase to facilitate handling, and reporter mRNAs are diluted to the desired concentration before injection. At the 1-cell stage, embryos are aligned on agarose injection plates and a defined volume of reporter mix is injected directly into the blastomere, followed by incubation at 28°C until the chosen collection time. Embryos are then pooled per replicate, flash-frozen, and stored at -80°C. This protocol yields injected, developmentally staged embryos that can be lysed and carried forward into the NaP-TRAP pulldown and downstream RNA analysis.

#### Materials

5X Pronase (see **REAGENTS AND SOLUTIONS**)

Methylene Blue Water (see **REAGENTS AND SOLUTIONS**)

Adult male and female zebrafish (*Danio rerio*)

*Adult zebrafish can be obtained from international stock centers such as the Zebrafish International Resource Center (ZIRC) or the European Zebrafish Resource Center (EZRC)*.

Embryo micro-injection Plate Mold (WPI, Model No. Z-MOLDS)

Microscope Calibration Slide (AmScope, Cat. No. MR095)

Microscopy for embryo micro-injection (ZEISS, Model No. Stemi 508)

Incubator for zebrafish embryo (VWR, Model No. 89511-422)

Micro-Injection Pump (WPI, Model No. PU830)

#### Zebrafish crossing, embryo collection, and injection

1. The evening prior to injections, prepare at least eight pairs of zebrafish for breeding in designated crossing tanks, selecting healthy males and females aged between 6 and 15 months.

*Zebrafish require a recovery period of 10-12 days before they can be crossed again*.

2. Prepare injection plates with 2% agarose gel in blue system water and an injection plate mold. Also prepare 2% agarose-coated gel plates for dechorionated embryo transfer.
3. The next morning when the light cycle has started, replace the water in the crossing tanks with fresh system water.
4. Once the water is changed, prime the fish with white light using a flashlight, remove the dividers, and wait for 15 minutes.
5. During this time, thaw mRNA reporter samples and dilute them to 20 ng/µL. Calibrate your micro-injection needle to have 0.5 nL equivalent bubble using 1 mm calibration slide (∼0.1mm). A total of 1nL, or two puffs, is used per embryo.

*The amount of mRNA micro-injected and number of embryos collected can be optimized based on the complexity of the library. For the 5’-UTR library of 10,088 reporters, we found that micro-injection of 20 pg and collection of 50 embryos per sample to be sufficient for building NaP-TRAP-sequencing library with a number of PCR cycles that limit PCR duplicates and for providing a robust representation of each reporter. For NaP-TRAP-qPCR experiments, reporters can be pooled and micro-injected together with the dsRed spike-in. For example, for the miR-430 experiment (Fig. 1E), we microinjected 20 pg of the miR-430 WT, 20 pg of the miR-430 mutant, and 120 pg of the dsRed reporter. We also collected 25 embryos per replicate instead of 50 given the much lower reporter complexity*.

6. After 15 minutes, collect all the embryos from crossing tanks using a plastic sieve into a 100 mm petri dish.
7. Dechorionate embryos:
  a. Dilute the 5x pronase to 1x using methylene blue water and keep the tube at 28°C.
  b. Transfer the embryos in a 35 mm dish, removing as much water as possible.
  c. Add 3 mL of 1X Pronase to the embryos, and incubate for 2-3 minutes at room temperature (or as soon as five consecutive chorions break when gently touched with forceps).
  d. Post incubation, transfer embryos in a 250 ml beaker filled with methylene blue water.
  e. Once the embryos settle at the bottom of the beaker, remove as much water as possible without losing embryos.
  f. Rigorously pour ∼250 ml of methylene blue water into the beaker directly onto the embryos. The water pressure will burst the chorion.
  g. Create a swirl with a glass pasteur pipette, and carefully collect dechorionated embryos and line them on an injection plate.

*Keep dechorionated embryos on agar-coated plates at all times to avoid cell lysis*.

8. Micro-inject 1 nL of mRNA reporter mix into the blastomere of 1-cell stage embryos.
9. Transfer the embryos to an agarose coated plate and incubate at 28°C until collection.

*During the incubation period, periodically check embryos for growth and remove the dead embryos*.

#### Embryo processing and storage

10. At collection time, collect embryos (25-50 depending on the type of experiment) in a 1.5 mL tube for each replicate, remove as much excess water as possible, and flash-freeze the embryos in liquid nitrogen.
11. After flash-freezing, embryos can be stored at -80°C for an extended period of time.
12. When ready to start NaP-TRAP, thaw the embryo tubes briefly on ice.
13. Add 500 µL of cold Lysis Buffer, and pipette mix the embryos.
14. Vortex lysate briefly then proceed to the **Basic Protocol 3** section.

#### Alternate Protocol 1: NaP-TRAP DELIVERY BY TRANSFECTION IN CULTURED MAMMALIAN CELLS

This alternative protocol describes how to deliver NaP-TRAP reporter mRNAs (individual reporters or complex libraries) into cultured mammalian cells by lipid-mediated transfection. Cells are seeded to reach the appropriate confluency at the time of transfection, and reporter mRNAs are mixed with a transfection reagent in serum-free medium to form mRNA–lipid complexes. Complexes are then added to cells and incubated under standard culture conditions to allow rapid cytoplasmic delivery and initiation of translation. At the desired time point, cells are washed, harvested, and flash-frozen or lysed directly for NaP-TRAP immunocapture and downstream RNA quantification. This procedure yields efficient, reproducible reporter delivery with robust FLAG-GFP translation, enabling NaP-TRAP measurements of cis-element effects in a mammalian cell context.

#### Additional Materials

Cultured cell lines

*We have used NaP-TRAP on HEK293T (RRID: CVCL_0063, ATCC Cat. No. CRL-3216) but successfully performed it in various other cell lines including MCF7 (RRID: CVCL_0031, ATCC Cat. No. HTB-22) and H9 (RRID:CVCL_1240, ATCC Cat. No. HTB-176)*.

*Cell lines should be periodically tested to ensure that cells are free from mycoplasma*.

DMEM complete medium

Opti-MEM I Reduced Serum Medium (Thermo Fisher Scientific, Cat. No. 31985062)

Gemcell SuperCalf Serum (Gemini, Cat. No. 100-510)

Dulbecco’s phosphate-buffered saline (1X DPBS) (Thermo Fisher Scientific, Cat. No. 14040-133)

Trypsin-Ethylenediaminetetraacetic acid (EDTA) (Thermo Fisher Scientific, Cat. No. 25200056)

Lipofectamine™ MessengerMAX™ (Thermo Fisher Scientific, Cat No. LMRNA001)

Cycloheximide from microbial, ≥94% (TLC) (Sigma-Aldrich, Cat. No. C7698) (see **REAGENTS AND SOLUTIONS**)

Lysis buffer (see **REAGENTS AND SOLUTIONS**)

DPBS + cycloheximide (see **REAGENTS AND SOLUTIONS**)

Serological pipettes, sterile, 25 mL, 10 mL and 5 mL (USA Scientific, Cat. Nos. 1072-5410, 1071-0810, 1071-0110)

Tubes 15 mL and 50 mL (VWR, Cat. No. 89039-658 and 89039-666)

6-well TC plate (USA Scientific, Cat. No. 087717)

Disposable PES Filter Units (Fisher Scientific, Cat. No. FB12566500)

Petri Dishes (Fisher Scientific, Cat. No. FB0875713)

CO2 Incubator (Thermo Fisher Scientific, Model No. HERACELL 150i)

Cell Counter (Thermo Fisher Scientific, Model No. AMQAX 1000)

Centrifuge for cell culture (Thermo Fisher Scientific, Model No. Cl2)

Microscopy for cell culture (ZEISS, Model No. Axiovert 25)

#### Cell culturing and transfection

1. Day 1: A night before transfection, seed around one million HEK293T cells in one well of a six well plate using DMEM complete media for each replicate. A minimum of three replicates should be anticipated per sample, *i*.*e*., 3 wells.
2. Incubate the cells overnight at 37°C in a humidified 5% CO2 incubator.
3. Day 2: The next day (morning or evening depending on incubation time before harvesting), transfect cells using Lipofectamine MessengerMAX according to manufacturer protocol. We use the following amounts:

Tube-1: 3 µL Lipofectamine Reagent plus Opti-MEM to total volume of 125 µL

Tube-2: 1-2 ug of *in vitro* transcribed RNA plus Opti-MEM to total volume of 125 µL

*This protocol is optimized for HEK293T cells, and adjustments may be required for different cell types. The ratio of Lipofectamine Reagent to mRNA reporters can be optimized using fluorescent reporters (e*.*g*., *GFP or DsRed) to ensure high transfection efficiency. Transfection reagents should be scaled according to manufacturer recommendations for different plate sizes*.

4. Vortex each tube separately at a low level, then incubate at RT for 10 min.
5. After incubating, add the diluted mRNA from Tube-2 to Opti-MEM-Lipofectamine mix in Tube-1, flick the tube to mix.
6. Incubate the mixture at RT for 5 minutes.
7. While the reaction mix is incubating, replace HEK293T cells media with 1 mL of fresh and prewarmed DMEM complete media.
8. Add the 250 µL of transfection mixture on top of the media drop by drop without disturbing the cells.

*Add the transfection mix gently on the surface of the media to prevent the cells from dislodging from the surface*.

9. Incubate the cells at 37°C in a humidified 5% CO2 incubator for 1 hour.
10. Post one hour incubation, wash the cells with 1x PBS and feed the cells with 2 mL of fresh prewarmed DMEM complete media.
11. Incubate the cells for an additional 5 hours before harvesting them for processing.

*Incubation can be extended to 12 hours if a lower density is seeded the day before transfection (e*.*g. 200-300k cells per well)*.

#### Cell processing and storage

12. After incubation, remove the plate from the incubator and add cycloheximide (100 mg/mL stock) to the culture medium to a final concentration of 100 µg/mL.
13. Incubate for 10 minutes at 37°C.
14. During this incubation, prepare 500 µL per sample of Lysis Buffer.

*Lysis Buffer can be prepared earlier and kept on ice*.

14. Move the plate on ice and aspirate the media.
15. Wash the cells twice with 1 mL of ice cold DPBS + cycloheximide (100 µg/mL).

*While washing the cells, make sure the cells do not detach during the wash step*.

16. Lyse the cells with 500 µL/well of Lysis Buffer.
17. Use a cell scraper to mechanically lift the cells off of the plate. Use a P1000 to pipette up and down to break up clumps.
18. Transfer the lysate from each well to a 1.5 mL low bind tube.
19. Incubate the tubes for 10 minutes on ice.
20. Proceed directly to **Basic Protocol 3**, or freeze lysate at -80°C.

#### Basic Protocol 3: NaP-TRAP PULLDOWN and RNA EXTRACTION

This protocol describes how to immunocapture actively translated NaP-TRAP reporters and prepare RNA for downstream quantification. Injected zebrafish embryos (or transfected cells) are lysed under conditions that preserve ribosome-nascent-chain complexes, and an aliquot of lysate is retained as the input fraction. Translating reporters are then immunocaptured via the FLAG tag on nascent-peptides using anti-FLAG magnetic beads, enriching ribosome-associated reporter mRNAs in the pulldown fraction. RNA is extracted in parallel from input and pulldown samples (with optional addition of spike-in RNAs during extraction for normalization). This workflow yields high-quality RNA from matched input and pulldown fractions, with robust enrichment of translated reporters in the pulldown, enabling reliable qPCR readouts and/or preparation for sequencing-based NaP-TRAP analyses.

#### Additional Materials

Lysis buffer (see **REAGENTS AND SOLUTIONS**)

Bead Wash Buffer (see **REAGENTS AND SOLUTIONS**)

Cycloheximide solution (see **REAGENTS AND SOLUTIONS**)

NaP-TRAP Wash Buffer (see **REAGENTS AND SOLUTIONS**)

TRIzol (Thermofisher Scientific, Cat. No. 15596026)

Spike-in mastermix (see **REAGENTS AND SOLUTIONS**)

TRI reagent (Thermo Fisher Scientific, Cat. No. 15596018)

GlycoBlue (Thermo Fisher Scientific, Cat. No. AM9516)

Pierce™ Anti-DYKDDDDK Magnetic Agarose (ThermoFisher Scientific, Cat. No. A36798)

Ethanol absolute (Sigma-Aldrich, Cat. No. 32205)

Chloroform (Sigma-Aldrich, Cat. No. 366927)

Isopropanol (American Bio, Cat. No. AB07015-01000)

Tube Rotator (Fisher Scientific, Cat. No. 11-676-341)

Biological safety cabinet (NUAIRE, Model No. NU-543-400)

3 mL Syringe with 25G needle (*e*.*g*., Fisher Scientific, Cat. No. 14-817-131)

Magnetic stand (Thermo Fisher Scientific, Cat. No. 12321D)

#### NaP-TRAP Pulldown

1. If frozen, thaw cell lysates on ice followed by gently vortexing. Next, triturate the mammalian or embryonic cell lysates kept on ice using a 25G syringe 5-10 times.

*Triturate each replicate and sample the same number of times to reduce variability between replicates. Avoid generating froth*.

2. Spin the tube at 16,000 xg for 5 minutes at 4°C.
3. Transfer the supernatant to a fresh 1.5 mL tube. If the total volume is less than 500 µL, bring it to 500 µL with Lysis Buffer.
4. Add 2 µL of DNase I in the tube and mix well with P1000.
5. Incubate the samples on ice for 15 minutes.
6. Add 500 µL of Lysis Buffer to the tube, mix well by pipette.
7. Remove 50 µL of this to a new 1.5 mL tube to act as the Input. The remaining sample is the Pulldown.
8. Store the Input at 4°C until the TRIzol addition step.

*This 50 µL will serve as input and is essential in the calculation of translation values (Fig. 1B)*.

9. Prepare magnetic beads:
  a. While the lysate is incubating, briefly vortex the anti-FLAG beads bottle to have homogenous distribution of beads.
  b. Transfer 20 µL of anti-FLAG beads per sample in a 1.5 mL low binding tube.
  c. Wash beads with 800 µL of Bead Wash Buffer for a total of three times by vortexing and using a magnetic stand.
  d. Keep beads in the last wash buffer on ice until ready to use.
10. Place prepared beads on the magnetic stand, and remove the supernatant. Transfer the Pulldown sample to the beads. Remove the tube from the magnet and pipette to mix.
11. Incubate the lysate-beads mix for 2 hours at 4°C on a Tube Rotator.
12. Towards the end of this incubation, prepare TRIzol with spike-in by adding 2 µL of prepared spike-in mastermix to 1 mL of TRIzol per sample. Vortex vigorously for 1-2 min. Keep on ice until Pulldown washing is complete.
13. Wash the Pulldown sample:
  a. Place the pulldown sample on the magnetic stand for ∼30 seconds at room temperature.
  b. Remove the supernatant.
  c. Add 800 µL of NaP-TRAP Wash Buffer.
  d. Vortex the sample tube for 15 seconds.
  e. Repeat the wash for a total of 3 times.
14. At the last wash step, separate the beads on the magnetic stand and remove the Wash Buffer.
15. Add 1 mL of TRIzol with spike-in to Pulldown samples.
16. Add 1 mL of TRIzol with spike-in Input samples.
17. For experiments with a sequencing readout, we recommend supplementing the TRIzol with spike-ins. The amount of spike-ins used might require adjustment depending on the number of embryos or cells and amount of NaP-TRAP reporters micro-injected or transfected (see **COMMENTARY** for more information about spike-in design).
18. Store -80°C or continue to the next steps.

#### RNA Extraction

19. If frozen, thaw the samples on ice.
20. To each thawed sample, add 200 µL of chloroform per 1 mL of TRIzol reagent.
21. Vortex for 1 min and incubate for 2 minutes at room temperature.
22. Centrifuge samples at 12,000 xg for 15 min at 4°C.
23. While tubes are spinning, add 2 µL of GlycoBlue in fresh 1.5 mL low binding tubes and label for each sample. The addition of GlycoBlue will help to precipitate the RNA and visualize the pellet, especially in pulldown samples.
24. Transfer 450-500 µL of aqueous upper phase containing the RNA to the new tube with Glycoblue. Be careful to avoid pipetting the interface layer containing genomic DNA. Transfer the same amount per sample to reduce noise.
25. Add 500 µL of (1 volume) of isopropanol and vortex for 30 seconds.
26. Incubate at -80°C for 10 min.
27. Centrifuge at 12,000 xg for 30 minutes at 4°C with tube hinges facing outside.
28. Carefully, remove the supernatant without disturbing the pellet. A blue translucent pellet should be visible. Avoid touching the pellet during pipetting to reduce the likelihood of losing the pellet.
29. Wash the pellet by adding 1 mL of ice-cold 75% ethanol. Ensure the pellet dislodges from the tube wall. Spin 5 min 12,000 xg. Repeat the wash once.
30. Remove ethanol and spin the tube at maximum speed for one minute.
31. Remove remaining ethanol using 20 µL pipette and dry the RNA pellets by leaving the tubes open for 5-10 min.

*Do not over dry the pellet as it will become difficult to resuspend, but it is critical to minimize EtOH presence before reverse transcription*.

32. Add 11 µL of Nuclease-Free Water and let the pellet incubate in water for 5 minutes on ice.
33. Pipette mix the water with the pellet, briefly vortex the mixture to resuspend RNA and quick spin with table top centrifuge.
34. RNA can be stored for ≤ 1 month at -20°C or long term at -80°C.
35. For high throughput sequencing read-out, continue to **Basic Protocol 4**. For RT-qPCR, continue to **Alternate Protocol 2**.

#### Basic Protocol 4: PREPARATION OF NaP-TRAP-SEQUENCING LIBRARIES

This protocol describes how to generate barcoded cDNA samples from matched Input and Pulldown RNA samples and convert them into Illumina-ready sequencing libraries. RNA samples are first reverse-transcribed using custom barcoded RT primers. Barcoded cDNA replicates are then pooled (keeping Input and Pulldown separate) and purified with AMPure beads to remove excess primers. Next, the minimal number of PCR cycles required for library amplification is determined using a small qPCR-based optimization. Libraries are then amplified with indexed Illumina primers, purified again, and quantified to enable equimolar pooling. If sequencing is performed in-house, libraries are denatured, diluted to the appropriate loading concentration, spiked with PhiX to increase sequence diversity, and loaded onto a MiSeq (or another Illumina platform). This workflow yields clean, appropriately sized libraries at sufficient concentration to cluster efficiently and produce reproducible read distributions for downstream NaP-TRAP-sequencing analysis.

#### Additional Materials

Superscript III enzyme (Thermo Fisher Scientific, Cat. No. 18080044)

RNase H (New England Biolabs, Cat. No. M0297S)

RNase If (New England Biolabs, Cat. No. M0243S)

D1000 Screen Tape and Sample Buffer (Agilent, Cat. No. 5067-5582, 5067-5583)

NEBNext Library Quant Kit for Illumina (NEB, Cat. No. E7630S)

AmpureXP Purification Beads (Beckman Coulter, Cat. No. A63880)

Illumina MiSeq Reagent Kit v2 (Illumina, Cat. No. MS-103-1001)

PhiX Control V3 (Illumina, Cat. No. FC-110-3001)

TapeStation System (Agilent, Model No. 4200)

PCR plate (BIO-RAD, Cat. No. MLL9601)

Adhesive PCR plate seal (BIO-RAD, Cat. No. MSB1001)

Real-Time PCR (BIO-RAD, Model No. CFX Opus 96)

MiSeq (Illumina, Model No. SY-410-1003)

#### Reverse Transcription

1. Thaw RNA if needed, then transfer 11 µL into a PCR tube
2. For each sample, transfer 1 µL of 10 mM dNTPs and 1 µL of 4 µM barcoded reverse transcription primers specific for the library in a PCR tube.

*The library specific reverse transcription primers carry a target RNA binding region, 6-nucleotide homemade barcodes, a 4-nucleotide anchor, 10-nucleotide UMI*, another *4-nucleotide anchor and part of the Illumina i7 adapter (Fig. 2E). We recommend using a different barcoded specific primer for each biological replicate allowing to pool all biological replicates together after reverse transcription (Fig. 2F). The cDNA primers are also the mix of two primers with similar binding sequence but are staggered by one base after the RNA binding region to create sequencing diversity. Recently, we have used a mixture of two primers with the same binding sequence but a differential number of untemplated nucleotides added immediately upstream the binding region for that purpose (see* ***COMMENTARY*** *for considerations for Illumina library sequencing preparation, and Support Fig. S1B)*.

3. Add the 11 µL of RNA sample to the PCR tube, mix by pipetting and gentle vortexing.
4. Denature RNA using the following thermocycler program:

2 min at 95°C

5 min at 65°C

5 min at 55°C

5. While samples incubate, prepare RTase mastermix:

4 µL 5X SSIII buffer

1 µL 100 mM DTT (from SSIII kit)

1 µL RNase out

1 µL SSIII

6. Once samples complete the 55°C incubation, quickly add 7 µL of RTase mastermix to each tube and pipette to mix. Spin the tubes briefly to bring all material to the bottom of the tube.
7. Return tubes to the thermocycler and incubate the RT reaction:
  a. If using a random hexamer primer, incubate samples for 10 min at 25°C followed by 55°C for 50 min.
  b. If using template-specific primers, skip the 25°C step and incubate at 55°C for one hour.
8. Deactivate the RTase at 70°C for 15 min.
9. Remove tubes from thermocycler to ice and add 1.5 µL each of RNase H and I per sample. Pipette to mix, briefly spin down, then return to the thermocycler and incubate at 37°C for 30 min.
10. Store cDNA at -20°C or continue to the next steps.

#### Library cDNA Purification

11. Pool all the barcoded cDNA replicates together in a 1.5 mL low binding tube (Fig. 2F).

*CRITICAL: Keep Input and Pulldown samples separate*.

*Pooling replicates at this stage decreases the number of samples to manage and ensures that each replicate undergoes the same level of amplification*.

12. Purify the library using AmpureXP beads.
  a. Prepare AmpureXP beads by vortexing to mix.
  b. Add 1.8x volume of AmpureXP beads into the sample tubes and pipette mix them using P200. *For consistency across all the samples, pipette mix the cDNA at least 20 times using P200*.
  c. Incubate the beads mix for 15 minutes at room temperature.
  d. Separate the beads by placing the tube on a magnetic stand and wait for 2 minutes.
  e. Remove supernatant.
  f. Wash the beads by adding freshly prepared 700 µL of 70% ethanol to remove contaminants.
  g. Wait for 10 seconds and remove the supernatant.
  h. Repeat wash for a total of two times. *Do not move the 1.5 mL tube off the magnetic stand during washes*.
  i. Air dry the beads to remove residual ethanol by opening the cap for 5 minutes. *Over drying of beads might impact the elution of the library*
  j. Add 30 µL of Nuclease free water.
  k. Resuspend the beads by vortex mix.
  l. Incubate the mix for 5 minutes at room temperature.
  m. Briefly spin the tube to collect the material at bottom.
  n. Keep the tube back on the magnetic stand and wait for 2 minutes.
  o. Once the beads are separated, collect the sample.
  p. To avoid bead carryover, leave around 1 µL volume.
  q. Store samples at -20°C or directly continue to amplification.

#### PCR Cycle Optimization for Library Amplification

13. To quantify the minimal number of cycles required for amplification. Set up a reaction as shown in below mentioned table in a 96 well plate on ice: 1 µL of cDNA from previous step 1 µL of I5_10nt_[id]_fwd 1 µL of I7_10nt_[id]_rev 12.5 µL of KAPA HiFi HotStart ReadyMix 9.5 µL of Nuclease-Free Water
14. Set up the qPCR reaction as mentioned below: 3 min at 95°C 35 cycles: 15 s at 98°C 20 s at 60°C 30 s at 72°C

*We recommend adding a melting curve as well to confirm the presence of a single amplicon when using a new primer pair*.

15. Calculate the minimal number of cycles required for library amplification by identifying the number of cycles at half of max relative fluorescence unit (RFU). Subtract 4 cycles from this amount to get the final number of cycles necessary.

*We notice that subtracting 4 cycles to the half of max relative fluorescence gives a number of cycles high enough to amplify the library to sufficient level for sequencing while limiting over-amplification leading to high number of PCR duplicates*.

#### Library Amplification and Purification

16. To amplify the cDNA library, prepare the following PCR reaction:

19 µL of cDNA from previous step

1.8 µL of I5_10nt_[id]_fwd

1.8 µL of I7_10nt_[id]_rev

30 µL of KAPA HiFi HotStart Polymerase Readymix

7.4 µL of Nuclease-Free Water

*Use a unique Illumina i5 and i7 index combination for each sample*.

17. Split the reaction in three PCR tubes, 20 µL in each tube.
18. Amplify the cDNA library as followed:

3 min at 95°C

**X cycles** (calculated in previous section): 15 s at 98°C

20 s at 60°C

30 s at 72°C

5 min at 72°C

Hold 12°C

*At the final extension step, the yield can be checked by D1000 TapeStation. Remove 1 µL from the sample, keeping the tubes in the thermocycler until confirmed. If the yield is too low, another 1-3 cycles can be added at this point*.

19. Purify the library using AMPure beads. Pool the three microtubes to 60 µL TV, then use 1.8X the amount of beads.
20. Elute library in 10 µL of Nuclease-Free Water. Store at -20°C or proceed to next steps.

*Library samples must be quantified prior to MiSeq sequencing. We recommend two different quantification methods depending on equipment availability and preference. One based on the NEB Library Quant Kit (Steps 21-27) and another one using a TapeStation (Steps 28-41)*.

#### Library Quantification by NEB Library Quant Kit

21. Thaw all the reagent on ice and pulse-vortex on a low setting.
22. Prepare all reagents as described in the kit protocol:
  a. Make a 1X dilution buffer with a 1:10 dilution of 10X NEBNext Library Quant Dilution Buffer in Nuclease-Free water.
  b. Add the Primer Mix to the Library Quant Mastermix tube and mix.
23. Using the Library Quant Dilution Buffer, make a library dilution of 1:1000, 1:10,000, 1:100,000 as specified below.
  a. Initial dilution: Make 1:1000 dilution of library sample in 1X dilution buffer by adding 999 µL of dilution buffer to 1 µL of library sample. Mix well by vortexing. *To properly mix the sample, add 1 µL of the library first in a 1*.*5 mL tube and then add the required volume of 1X dilution buffer*.
  b. Prepare second dilution: Add 10 µL of 1:1000 diluted library in a 1.5 mL tube and add 90 µL of 1X dilution buffer. Mix well by vortexing.
  c. Final dilution: To make 1:100,000 dilution, add 10 µL of 1:10,000 diluted library in a 1.5 mL tube and add 90 µL of 1X dilution buffer. Mix well by vortexing.
  d. Use only the 1:10,000 and 1:100,000 dilutions for library quantification.
24. Prepare a 96-well plate by adding 16 µL of enzyme mix to each well. Plan to do each dilution in triplicate. Add 4 µL of respective standard or library dilutions to each well. Also prepare a no template control (NTC) reaction. Pipette each well at least 5 times to mix while minimizing bubbles.
25. Run a qPCR assay with following program: 1 min at 95°C 35 cycles: 15 s at 95°C 10 s at 63°C

26. Use the NEB qPCR BioCalculator to calculate library concentrations using the standard curve and Cq values: https://nebiocalculator.neb.com/#!/qPCRGen
27. To pool multiple libraries into one final pool, calculate the volume of each library necessary to achieve a 1:1 molar ratio using C1V1=C2V2. To run on a MiSeq sequencer, we typically make a 20 nM pool; with 2 libraries, this means you should add 10 nM of each library. Repeat the quantification kit with the pooled library to ensure the final concentration is as expected.

#### Library Quantification by TapeStation

28. Thaw D1000 sample buffer at RT for 30 minutes.
29. Add 3 µL of D1000 sample buffer in 200 µL Optical tubes
30. Add 1 µL of sample library and seal the tube with Optical strips.
31. In another tube add 1 µL of DNA ladder with 3 µL of D1000 sample buffer.

*An electronic ladder can also be added to measure amplicon length*.

32. Vortex mix the library and ladder samples for 1 minute.
33. Spin the tube to collect all the liquid at the bottom of the tube.
34. Start the TapeStation program.
35. Select the well where you aim to place your library sample.
36. Place a fresh D1000 Tape in the TapeStation instrument.
37. Load the dedicated tip box loaded with tips.
38. Remove the cap strip and load the sample tube in a dedicated well.

*Remove the cap strip carefully to avoid dislodging the sample*. .

39. Close the instrument lid and start the program.
40. Once the run is over, create a report as a PDF file and verify the amplicon length as well as concentration.
41. If multiple libraries are to be pooled, combine for equal molarity as described previously.

#### Sequencing on MiSeq

42. For cost-effectiveness, we sequence on a MiSeq Reagent Kit v2 flow cell, which yields enough reads to assess library homogeneity and the fraction mapping to spike-ins. Any Illumina platform or flow cell with similar or higher output can be used. Thaw the MiSeq Reagent Kit v2 cartridge by putting it at 4°C overnight.

*The cartridge can also be thawed on the day of the run by incubating the cartridge in a water bath at room temperature*.

43. Thaw HT1 buffer at room temperature and put on ice once thawed.
44. Once the library is quantified, make an aliquot and dilute it to 4 nM.
45. Prepare a 4 nM PhiX dilution.
46. Add 30% PhiX by adding 1.5 µL of 4 nM PhiX into 3.5 µL of the 4 nM library sample in a 1.5 mL tube. Vortex to mix and spin to bring all liquid to the bottom of the tube.

*The amount of PhiX should be adjusted based on the diversity of the library. For low diversity libraries, we have used up to 60% of PhiX. See* ***COMMENTARY*** *for more considerations about Illumina library preparation*.

47. Denature the sample by adding 5 µL of 200 mM NaOH. Vortex, spin down, and incubate at RT for 5 minutes.
48. Post incubation, add 990 µL of ice-cold HT1 buffer. At this step the final library concentration is 20 pM.
49. Dilute the library to 8 pM by adding HT 1 buffer, add 240 µL of 20 pM library and 360 µL of HT1 buffer. Mix the library by inversion and keep it on ice.

*Different libraries might need optimization for final loading concentration on MiSeq for maximum clustering on the flow cell*.

50. Prepare the MiSeq system by completing required washes and uploading the sample sheet.
51. Follow the instructions on the MiSeq sequencer and load the sample and cartridge. Ensure the MiSeq program is set to save the results to BaseSpace for easy access.
52. Once the run is complete, complete required washes and move on to **Basic Protocol 5**..

#### Alternate Protocol 2: NaP-TRAP-qPCR MODULE FOR LOW-COST VALIDATION

This alternate protocol describes a streamlined NaP-TRAP-qPCR module for rapid, low-cost validation of translational regulation using individual or small pools of NaP-TRAP reporters. Following NaP-TRAP pulldown and parallel RNA extraction from matched Input and Pulldown fractions, cDNA is synthesized and reporter abundance is quantified by qPCR using primer sets targeting each reporter. Enrichment of each reporter in the Pulldown relative to Input provides a quantitative readout of translation, and comparisons across reporters (and/or conditions) reveal the impact of specific 5′-UTR elements on translation. This module yields robust, reproducible Pulldown/Input enrichment measurements without sequencing, enabling efficient troubleshooting, smaller scale experiments, and focused validation prior to scaling to NaP-TRAP– sequencing.

#### Additional Materials

Sso Advanced Universal SYBR Green Supermix (Biorad, Cat. No. 1725270) Reporter-specific qPCR oligos (see **Supporting Information**: Table S1) Laptop or desktop for data analysis

#### Reverse Transcription

1. Thaw RNA if needed, then transfer 11 µL into a PCR tube. For each 11 µL sample, transfer 1 µL of 10 mM dNTPs and 1 µL of 50ng/µL random hexamer primers to a PCR tube.
2. Heat the mixture at 65°C for 5 min, then incubate on ice for 1 min.
3. Prepare the following mastermix: 4 µL 5X First-Strand Buffer 1 µL 0.1 M DTT 1 µL RNaseOUT 1 µL SuperScript III RT
4. Mix by pipetting up and down. Incubate for the following program: 5 min at 25°C 1 hr at 50°C 15 min at 70°C
5. Store cDNA at -20°C or continue to the next steps.

#### qPCR

6. Bring the total RT reaction volume to 100 µL (+80 µL). Then create a 1:5 dilution to use for qPCR.
7. Plan your plate setup and total reactions to calculate mastermix volumes. For two reporters, there will be two experimental primer sets plus a dsRed primer set. We recommend performing 3 technical replicates per sample (Fig. 2C).
8. Mix the following qPCR components per mastermix (make one mastermix per primer set): 7.5 µL Sso advanced universal SYBR 2X MM 2 µL 10 uM primer set* 2.5 µL Nuclease-Free Water *Primer Set 1: 1 µL each of reporter 1 forward with common reporter reverse (*e*.*g*., no_miR430_qPCR_ fwd with T7_qPCR _rev). Primer Set 2: 1 µL each of reporter 2 forward with common reporter reverse (*e*.*g*., or miR430_qPCR_ fwd with T7_qPCR _rev). Primer Set 3: 1 µL each of dsred_qpcr_fwd and dsred_qpcr_rev. *The number of primer sets depends on the number of NaP-TRAP reporters pooled per sample. In this case the reverse primer is the common one, but it could be the forward primer if the reverse primer is used to discriminate between reporters (see* ***COMMENTARY*** *for more information about reporter design for a qPCR readout)*.

9. Add 12 µL of mastermix to each well of a 96-well PCR.
10. Add 2 µL of each 1:5 diluted sample.
11. If needed, spin down the plate to ensure all liquid is at the bottom of the plate.
12. Run the qPCR with the standard SsoAdvanced SYBR protocol, and include the melt curve:

3 min at 95°C

39 cycles: 15 s at 95°C

20 s at 60°C

Melt curve: 65°C-95°C, 0.5°C and 5 s/cycle

13. For each reporter, the NaP-TRAP-qPCR translation value is calculated using the 2^-(ΔΔCq)^ approach. C_q_ values of technical replicates are first averaged for each sample. Then, pulldown and input averaged C_q_ values are first normalized by the dsRed spike-in (first ΔC_q_). The ΔC_q_ of the pulldown is normalized by the ΔC_q_ of the input to get the ΔΔC_q_. To measure reporter abundance changes between two time points the ΔC_q_ of input time point 1 can be normalized by ΔC_q_ of input time point 2 to get the ΔΔC_q_ (see **Supporting Information**: Table S2).

#### Basic Protocol 5: COMPUTATIONAL ANALYSIS OF NaP-TRAP MPRA DATA

This protocol describes how to process NaP-TRAP-sequencing data to quantify reporter translation at scale and generate publication-ready outputs. Starting from raw FASTQ files for matched Input and Pulldown samples, reads are trimmed, demultiplexed, aligned to their corresponding reporters, and summarized into count tables. Counts are then normalized (optionally using spike-in reporters) and reporter translation calculated as Pulldown/Input enrichment. This workflow includes QC analyses to assess library complexity, read distribution, and replicate correlation. Finally, this analysis includes a kmer enrichment analysis for reporters with high and low translation. This computational pipeline yields reporter-level translation values and accompanying tables/plots that can be used for downstream statistical testing, element ranking, and interpretation of *cis*-regulatory effects.

The primary goals of the first few steps (**Data processing - Read trimming, UMI capture and demultiplexing** and **Data processing - Bowtie aligning**) is to extract replicate reads by demultiplexing FASTQ files using the custom barcodes added during reverse transcription and aligning the demultiplexed reads to the library to generate .`sam/.bam` alignment files (Fig. 4). Alternative pipelines that achieve the same result can also be employed, and the resulting .`sam/.bam` alignment files can be directly utilized for counting (**Data processing - Counting**). We normally use the `LabxDB` (Vejnar and Giraldez, 2020) and the `lxpipe` pipeline (https://github.com/vejnar/LabxPipe) to store and annotate FASTQ files and perform read trimming, UMI capture, demultiplexing and aligning at scale and with ease. However, here we provide a step-by-step procedure to achieve these steps manually using `ReadKnead` (https://github.com/vejnar/ReadKnead) and bowtie2 (Langmead and Salzberg, 2012) for the HEK293T and zebrafish NaP-TRAP experiments from Strayer *et al*. (Strayer et al., 2024) that can be downloaded from the Sequence Read Archive (SRA). Specifically, HEK293T input (SRR31519530), HEK293T pulldown (SRR31519545), zebrafish 2h input (SRR35883406), zebrafish 2h pulldown (SRR35883405), zebrafish 6h input (SRR35883403), and zebrafish 6h pulldown (SRR35883402) (see **Supporting Information**: Table S3 for more information for each SRA sample). For zebrafish samples, we only analyze hard-encoded poly(A) (pA) datasets and omit sv40 datasets for simplicity.

#### Materials

`sra-tools` (https://trace.ncbi.nlm.nih.gov/Traces/sra/sra.cgi?view=software, SRA Toolkit Development Team)

`ReadKnead` (https://github.com/vejnar/ReadKnead)

`bowtie2`^55^

`zstandard` (https://github.com/facebook/zstd)

`samtools`^56^

`python` (https://www.python.org)

`git` (https://git-scm.com)

NaP-TRAP protocol GitHub^57^ (https://github.com/beaudoinlab/nap_trap_protocol)

Workstation or HPC for running data analysis

#### Data processing - Read trimming, UMI capture and demultiplexing

1. Set up a project directory. `mkdir nap_trap`
2. Make a new directory to store downloaded FASTQ files. `cd nap_trap mkdir sra_fastq_files cd sra_fastq_files`
3. Download the FASTQ files. Here we show an example with the HEK293T NaP-TRAP **Input** (SRR31519530). These files can be retrieved using the **SRA Toolkit (**`sra-tools`), but any alternative approach that yields the corresponding FASTQ files (e.g., direct download from SRA) also works. `sra-tools` can be installed in a conda environment. `conda install -n env_name -c conda-forge -c bioconda sra-tools conda activate env_name prefetch --output-directory . SRR31519530 fasterq-dump SRR31519530`

*Repeat* `prefetch` *and* `fasterq-dump` *for SRR31519545, SRR35883402, SRR35883403, SRR35883405, and SRR35883406. This should output FASTQ files for read 1 and read 2 of each sample in* `sra_fastq_files`.

4. Make new directories for demultiplexing and a subfolder for each sample.

~~~
   cd ..
   mkdir sra_demultiplexing
   cd sra_demultiplexing
   mkdir hek293_input
   mkdir hek293_pulldown
   mkdir 2h_input
   mkdir 2h_pulldown
   mkdir 6h_input
   mkdir 6h_pulldown
~~~

5. Define operations to be performed on read 1 and read 2 for each sample in individual JSON files (see examples `SRR31519530_r1_ops.json` and `SRR31519530_r2_ops.json` for sample HEK293T input). These files and the ones for HEK293T pulldown and zebrafish 2h input, 2h pulldown, 6h input and 6h pulldown are also available on our GitHub (Madhavan and Strayer, 2025) in `/supplementary/demultiplexing/.` Save `JSON` files in `sra_demultiplexing`.

*These operations include sample naming, read trimming, UMI clipping, and barcode demultiplexing. Briefly, Read 1 is trimmed using the fixed sequence immediately downstream of the insert. Read 2 is trimmed using the binding region of the RT primer, demultiplexed using homemade barcode, trimmed for the fix 4-nucleotides anchor sequence, and clipped for the 10-nucleotide UMI (added to the read name following “#”) (see Fig. 2E). If a fixed sequence is present upstream of the insert, an extra trimming step at the 5’ end of read 1 can be added (Fig. 2E)*.

6. Run `ReadKnead` with the following command:

~~~
readknead -fq_fnames_r1 “../sra_fastq_files/SRR31519530_1.fastq”\
       -fq_fnames_r2 “../sra_fastq_files/SRR31519530_2.fastq” \
       -fq_path_out “hek293_input” \
       -fq_fname_out_r1 “SRR31519530_[DPX]_1.fastq.zst” \
       -fq_fname_out_r2 “SRR31519530_[DPX]_2.fastq.zst” \
       -fq_command_out “zstd,-,-fo” \
       -ops_r1_path “SRR31519530_r1_ops.json” \
       -ops_r2_path “SRR31519530_r2_ops.json” \
       -report_path “hek293_input/preparing_report.json” \
       -label “hek293_input” \
       -ascii_min “33” \
       -stats_in_path “hek293_input/stats_in_hek293_input” \
       -stats_out_path “hek293_input/stats_out_hek293_input”
~~~

*This step can take a long time for large FASTQ files. In such a case, we suggest parallelizing* `ReadKnead` *using the* `-num_worker argument`. *The output demultiplexed FASTQ files are compressed using the* `Zstandard` (https://github.com/facebook/zstd) *to limit disk space usage. The* `[DPX]` *placeholder in file names is replaced by the corresponding homemade barcode sequence in the demultiplexed FASTQ output. A ReadKnead summary report is generated at* *hek293_input/preparing_report*.*json*, *which provides step-by-step processing statistics, including the number of reads assigned to each in-house barcode. Command to demultiplex the other samples is the same with appropriate changes for file names, label, and input and output paths. At the end of this step, each folder should contain demultiplexed FASTQ files for read 1 and read 2. Note that for zebrafish samples, it is normal to obtain a large fraction of undetermined reads during demultiplexing; these reads originate from two additional samples (sv40) that are not analyzed in this protocol*.

#### Data processing - Bowtie aligning

7. Make a fasta file of the NaP-TRAP library containing the insert name and sequence of each insert. For the NaP-TRAP 5’-UTR library it is composed of 11,088 inserts and 5 spike-ins and the fasta file can be found on GitHub (Madhavan and Strayer, 2025) in `/supplementary/bowtie/reporters.fa`.

*It is essential to add either #reporter or #spikein to the name of the library insert or spike-in sequences in the fasta file. This will be used by the program to identify spike-ins during normalization. The analysis pipeline will detect them and consider them accordingly, but will omit them for querying and outputting data. Make sure that no reporter name contains the string “spikein” otherwise it will be considered as a spike-in*.

*If the insert length is much longer than the sequencing read length, we suggest truncating the insert length to match read length. This is particularly important if you used a sliding window strategy with a small step to make the NaP-TRAP library (Fig. 2D). In such a case, shorter reads could map perfectly to more than 1 insert (e*.*g*., *starting a position 1 of the proper insert and position 25 of the next and wrong insert)*.

8. Build a `bowtie2` index from the NaP-TRAP library fasta file:

~~~
    cd ..
    mkdir bowtie2_index
    cd bowtie2_index
    bowtie2-build ../reporters.fa seq
~~~

*This creates an index using the library as the reference genome, with each insert acting as a separate chromosome*.

9. For each sample, ONLY align the Read 1 `fastq` file (only for files ending in “`_R1.fastq.zst`”) to the custom reference genome using `bowtie2`. Here is an example for sample SRR31519530_ACTTGA:

~~~
   cd ..
   mkdir aligning cd aligning
   bowtie2 -x “../bowtie2_index/seq” \
   -U
   “../sra_demultiplexing/hek293_input/SRR31519530_ACTTGA_R1.fastq.zst”\
   -S “SRR31519530_ACTTGA.sam” \
   --phred33 \
   --no-mixed \
   --no-discordant \
   --no-unal \
   --end-to-end
~~~

*Repeat for each sample, updating input and output paths accordingly. Successful completion of this step will generate one SAM file per demultiplexed sample (16 total) in the* `sra_demultiplexing` folder.

*Native support for* `Zstandard` *compressed FASTQ files was added in version 2*.*4*.*3 or newer of* `bowtie2`. *Make sure to NOT use an older version. Alternatively, use a different file compression method (e*.*g*., *gz or bz2) or uncompressed files*.

*While Read 2 was essential for demultiplexing and retrieving the UMI, Read 1 alone is adequate to map each read to a specific insert sequence. Consequently, all subsequent analyses depend exclusively on the examination of Read 1. While other aligners like* `STAR` *are available, we find* `bowtie2` *to be more effective for aligning reads to reference genomes that have numerous short chromosomes and lack splicing events*.

#### Data processing - Counting

10. Install the necessary scripts by cloning the nap_trap_protocol GitHub repository (Madhavan and Strayer, 2025) (https://github.com/beaudoinlab/nap_trap_protocol).

~~~
  cd ..
  mkdir scripts
  cd scripts
  git clone -b protocol \ https://github.com/beaudoinlab/nap_trap_protocol.git
  cd nap_trap_protocol
  pip install -e .
~~~

*Make sure that* `pip` *is updated to avoid compatibility errors (*`pip install --upgrade pip`*). We recommend running the pip command in a conda (or similar) environment containing the software requirements mentioned on the nap_trap_protocol GitHub* (Madhavan and Strayer, 2025), *including using python version 3*.*11 or above*.

11. Run `count` command to count reads mapping to individual reporters. The `count_reads` program is UMI aware and will report raw read counts after removal of PCR duplicates. Output is a `JSON` file containing a dictionary with the samples as keys and a dictionary of insert name : raw count as values.

~~~
   cd ../..
   mkdir counting
   cd counting
   naptrap count -i “/full_path_to_root_directory/aligning/*.sam” \
   -e “nap_trap_protocol” \
   -o $(pwd) \
   -t “path_to_tmp_folder” \
   -m1 100 \
   -d1 4 \
   -p 2
~~~

*The* `-i` *argument is a regular expression used to select all SAM files from the* `aligning` *directory (note that you must provide the full path to the root directory). Input alignment files can be in either* .*sam*, .*bam or* .*sam*.*zst* *format (samtools version 1*.*2 or above is required if using the* .*bam* *format)*. `-t` *is a temporary path where intermediary files will be written and deleted after completion of the program*. `-m1` *and* `-d1` *arguments must be used wisely depending on the experiment and context. They specify the minimum number of matches (determined from the CIGAR string) and maximum number of edit distance (determined by the NM tag) for a read alignment to be included in the counting. In this example, we required at least 100 matches and a maximum edit distance of 4 (the number of edits needed for a read to perfectly align with the reference sequence). These values will change depending on the quality of the oligo pool, the insert and read length as well as the flexibility in allowing the counting of non perfect sequences. We found that oligo pools ordered from leading companies like Twist Bioscience allow for using* `-d1 0` *and work only with reads perfectly matching their corresponding inserts. For large samples, the process can be run on multiple processors using the* `-p` *argument*.

12. Run the naptrap `read_cutoff` command to generate histograms displaying the raw read counts for each sample and to create a table that lists the number of reporters meeting different minimum read count thresholds (Fig. 4). These outputs are important for determining the suitable threshold for building the database, as explained in the following section.

~~~
   naptrap read_cutoff -i “name_of_count_file.json” \
   -o $(pwd)/ \
   -f “pdf”
~~~

*By default, the script generates a table (**Cutoff_vs_Count_table*.*csv**) for each sample listed in the JSON count file. To restrict the output to specific samples, use the* `-d` *option and provide a list of sample names matching the corresponding dictionary keys in the JSON file. Histograms are saved in the newly created* `figures` *folder (see Fig. 3A for an example)*.

**Figure 3.**
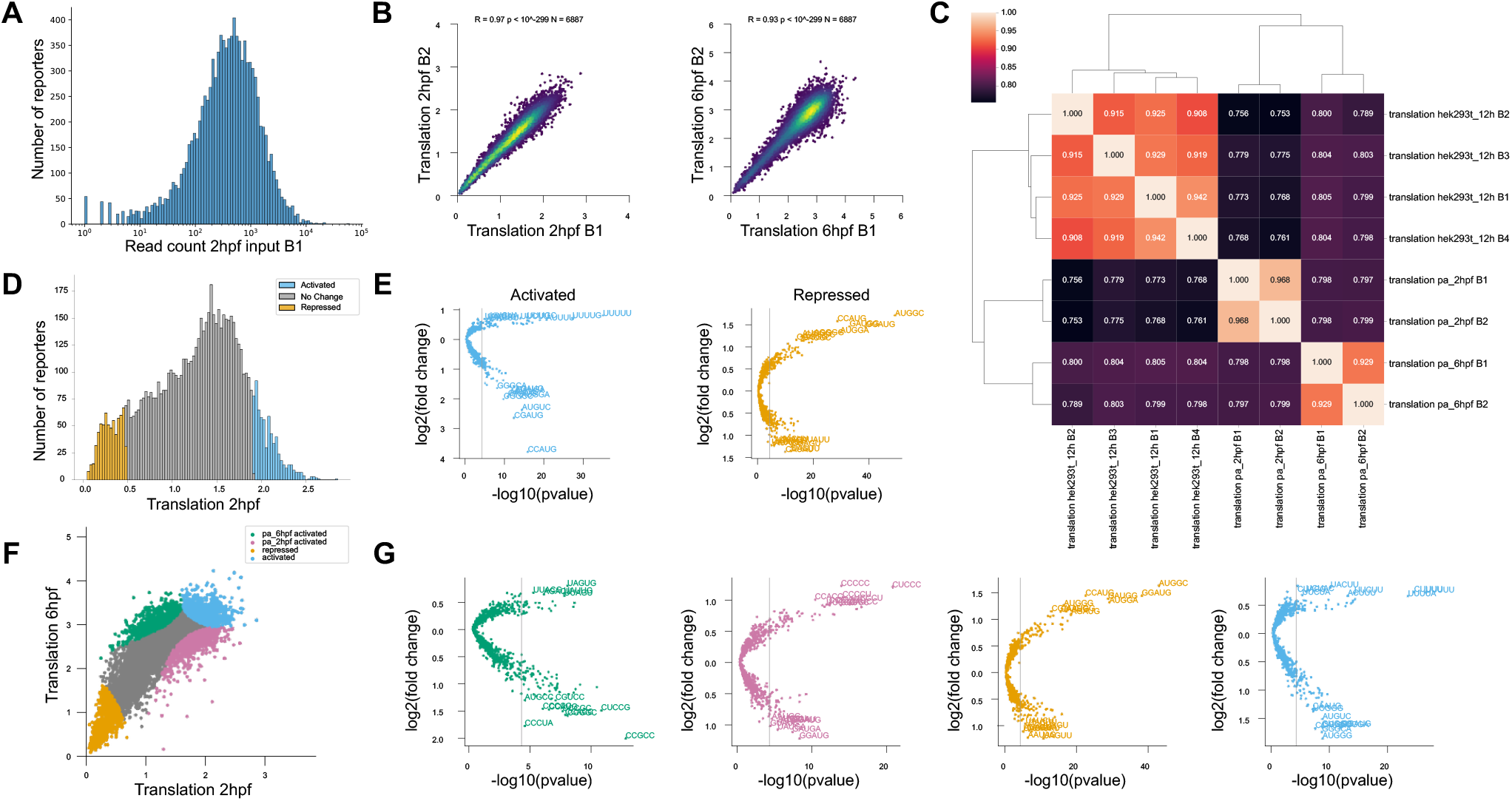
Expected results from NaP-TRAP MPRA analysis pipeline. (**A**) Example histogram of per-insert read counts for one Input replicate with 7,839,828 reads. (**B**) Representative replicate-to-replicate correlations of NaP-TRAP translation efficiency (TE; Pulldown/Input) at 2 hpf and 6 hpf (*R* values are Pearson correlations). (**C**) Hierarchical clustering heatmap of Pearson correlations showing TE similarity across replicates and conditions (zebrafish 2 hpf, zebrafish 6 hpf, and HEK293T). (**D**) Distribution of NaP-TRAP TE values at 2 hpf, with the top and bottom 10% of reporters highlighted as activated (blue) and repressed (orange), respectively. (**E**) Example k-mer enrichment analysis for activated and repressed reporter sets at 2 hpf. (**F**) Scatter plot comparing reporter NaP-TRAP TE at 2 versus 6 hpf, highlighting four reporter sets: reporters activated (blue) and repressed (orange) at both stages, reporters with higher TE at 2 hpf than at 6 hpf (2 hpf activated; green), and reporters with high TE at 6 hpf than at 2 hpf (6 hpf activated; pink). (**G**) k-mer enrichment analyses for each of the four reporter sets defined in (F): 2 hpf activated (green), 6 hpf activated (pink), globally repressed (orange) and globally activated (blue).

**Figure 4.**
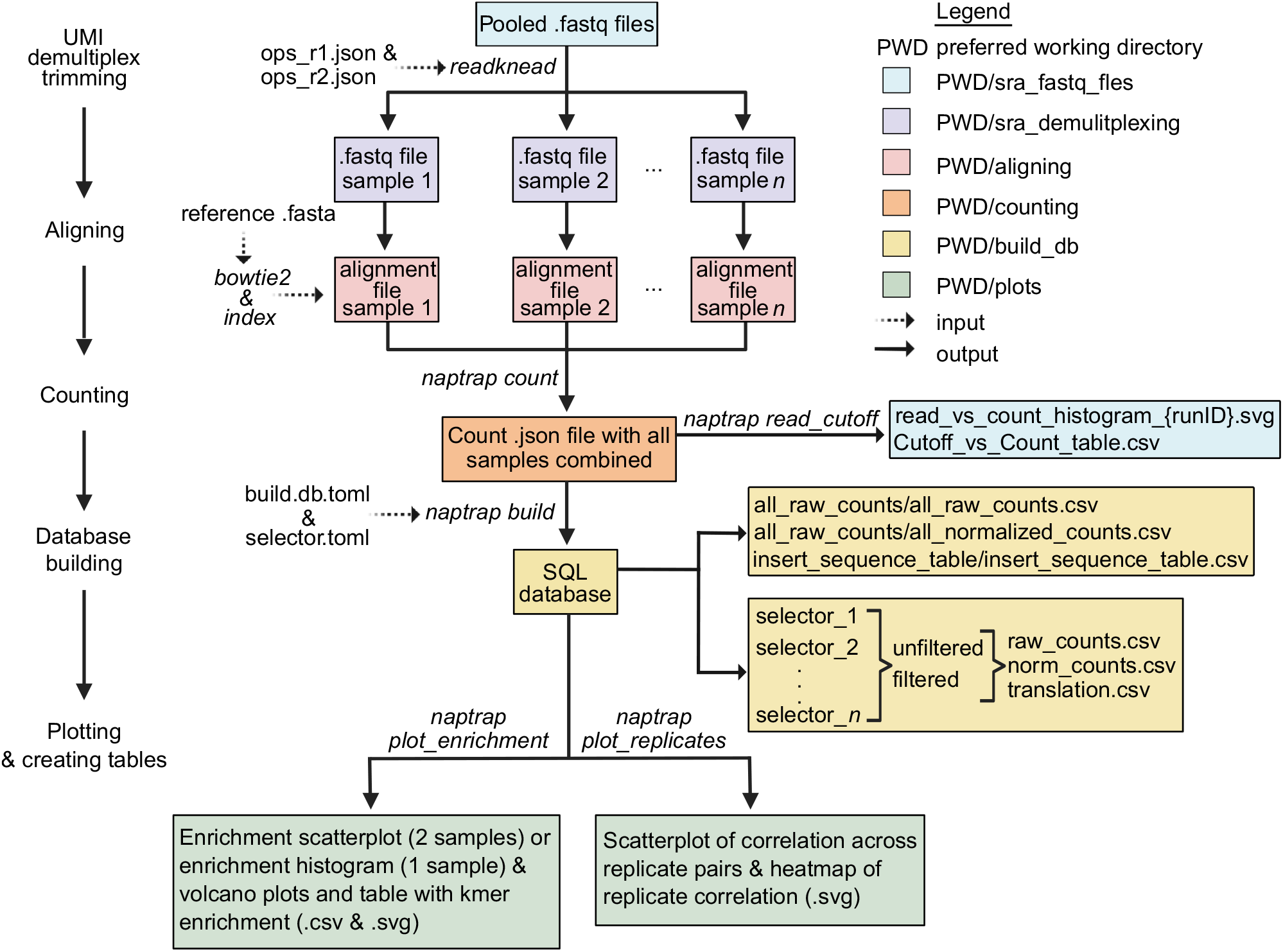
Computational pipeline for NaP-TRAP MPRA analysis. Pooled FASTQ files are first demultiplexed with UMI-aware adaptor trimming, then aligned to a user-generated bowtie2 index. The resulting alignment files are processed with the naptrap pipeline to generate a combined `.json` count file across all samples and to assess read-count cutoffs. Count data are then compiled into an SQL database using two user-edited `.toml` configuration files, enabling flexible sample selection and streamlined downstream analysis. The naptrap pipeline includes plotting commands to rapidly assess replicate correlation, visualize NaP-TRAP TE distributions (histograms and scatterplots), and identify enriched k-mers in reporter subgroups defined by TE. The workflow also generates various count and translation tables to facilitate downstream analyses.

#### Data processing - Building database and creating tables

13. Make a new directory for building the database and creating the tables.

~~~
   cd ..
   mkdir build_db
   cd build_db
~~~

14. Create and save the build_db.toml file in the build_db directory.

*The* *build_db*.*toml* *is where users enter crucial information about experimental configurations, paths to input files, sample details, features to extract and include in the database, analyses to conduct, and the function for calculating translation. An example of a* *build_db*.*toml* *file with extensive annotation and instructions can be found on GitHub* (Madhavan and Strayer, 2025) *at* *supplementary/builddb/build_db*.*toml*. *Instructions on how to complete the* *build_db*.*toml* *are also available in the GitHub readme*.

15. Create and save the selector.toml file in the `build_db` directory. Ensure that the path to `selector.toml` is specified in the `build_db.toml` file.

*The* *selector*.*toml* *file is where users define the samples and filtering parameters for various analyses. For instance, although our database contains samples from both fish and mammalian cell lines, we may choose to concentrate certain analyses solely on fish samples that meet the filtering parameters. We recommend setting a minimum threshold for raw reads across all input replicates, ensuring that only inserts with more than X number of reads in every replicate of all samples will pass the filter and be included in the analysis. In this analysis, we applied a threshold of 100 raw reads. Instructions for completing the* *selector*.*toml* *can be found in the GitHub* (Madhavan and Strayer, 2025) *readme*.

16. Run the `build` command to build the SQL database and generate tables for raw counts, normalized counts, translation, and average translation for all samples. Additionally, similar tables are produced for the different selectors outlined in the selector.toml for the unfiltered and filtered datasets, based on the specified minimum read thresholds (see Fig. 4).

`naptrap build --toml_path build_db.toml`

*Make sure that all necessary Python libraries are installed by checking the requirements on GitHub* (Madhavan and Strayer, 2025).

#### Data analysis - Replicate correlation and kmer enrichment

17. Make a new directory for generating plots.

~~~
   cd ..
   mkdir plots
   cd plots
~~~

18. Run the `plot_replicates` command to quickly assess the correlation of mean translation values among replicates. This script generates a heatmap of r-values between replicates as well as scatter plots to better visualize correlations between mean translation values (Fig. 3B,C and Fig. 4).

~~~
   naptrap plot_replicates \
   --db “../build_db/utr5.db” \
   --out $(pwd) \
   --selector “polyA_hek293t” \
   --fig_format “pdf”
~~~

19. Run the `plot_enrichment` command to analyze the kmer enrichment in different groups of reporters. When a single sample is given, the script creates a histogram of mean translation values and identifies two groups: the most repressed and activated reporters (Fig. 3D).

~~~
  naptrap plot_enrichment \
  --db “../build_db/utr5.db” \
  --out $(pwd) \
  --selector “polyA_fish” \
  --samples “pa_2hpf” \
  --klen 5 \
  --fig_format “pdf”
~~~

20. When two samples are provided, the scripts generate a scatter plot of mean translation values identifying four groups of reporters: i) repressed, ii) activated, iii) high in sample 1 but low in sample 2, and iv) low in sample 1 but high in sample 2 (Fig. 3F).

~~~
  naptrap plot_enrichment \
  --db “../build_db/utr5.db” \
  --out $(pwd) \
  --selector “polyA_fish” \
  --samples “pa_2hpf” “pa_6hpf” \
  --klen 5 \
  --fig_format ‘pdf
~~~

In both cases, the groups are determined based on the `rnum` value specified in the enrichment analysis section of the `build_db.toml` file (*e*.*g*., 0.1). Finally, the script produces kmer enrichment plots (Fig. 3E and G) and tables (Fig. 4) for each of the groups.

### REAGENTS AND SOLUTIONS

#### Agarose gel 1% (wt/vol)

Add 1 g of powder agarose to 100 mL of 1× TAE buffer and microwave until the agarose is molten. Allow the mixture to cool to ∼50°C, add 2 µL of EtBr. Pour into a gel forming apparatus and let sit for 30 min. Use immediately, or store in the dark at 4°C and use within 24 hrs.

#### Bead Wash Buffer

Prepare the reaction by combining 1× Salt Buffer and 2 mM DTT (Thermo Fisher Scientific, Cat. No. P2325) and bring to the desired final volume with nuclease-free water.

#### Cycloheximide solution (100 mg/mL)

Dissolve 0.1 g of cycloheximide powder (Sigma-Aldrich #01810-1G) in 1 mL molecular grade DMSO (Sigma Aldrich, Cat. No. D8418). Vortex to mix fully, then aliquot into 1.5 mL tubes and store at -80°C.

#### DMEM complete medium

Under a cell-culture hood, add 50 mL iron-supplemented fetal bovine serum for cell culture into 450 mL DMEM supplemented with high glucose (Thermo Fisher Scientific, Cat. No. 11965092) and L-Glutamine (Thermo Fisher Scientific, Cat. No. 25030081). Filter the mixture through a 0.2 µM filter unit. Store in the fridge for up to 1 month.

#### DPBS + cycloheximide

Resuspend cycloheximide powder (Sigma-Aldrich #01810-1G) to 100 µg/mL in Dulbecco’s phosphate-buffered saline (DPBS) (ThermoFisher Scientific #14190144). Aliquot into 1.5 mL tubes and store at -80°C.

#### Lysis Buffer

Prepare the buffer to a final composition of 1× Salt Buffer, 1% (v/v) Triton X-100 (Sigma, Cat. No. T8787-50ML), 2 mM DTT (Thermo Fisher Scientific, Cat. No. P2325), 1× Protease Inhibitor (from a 25× stock), 40 U/mL RNaseOUT (Thermo Fisher Scientific, Cat. No. 10777019), and 100 µg/mL cycloheximide (Sigma-Aldrich, Cat. No. 01810-1G). Bring to the desired final volume with Nuclease-Free Water.

#### Methylene Blue Water

Add 5 mL of methylene blue (Sigma, Cat. No. 122965-43-9) to 10L zebrafish facility system water.

#### NaP-TRAP Wash Buffer

Prepare a final concentration of 400 mM NaCl (Thermo Fisher Scientific, Cat. No. AM9760G) to total volume with Lysis Buffer.

#### PCR amplification primers

Dissolve PCR amplification primers to make 10 μM stock solutions in Nuclease-Free Water and stored at -20°C for at least 1 year.

#### 5X Pronase

Resuspend powder Pronase (Millipore Sigma, cat. No. P8811) to 5 mg/mL with deionized water.

#### 25X Protease Inhibitor

Dissolve 1 pellet of Protease Inhibitor (Sigma Aldrich, 11873580001) in 2 mL of Nuclease-Free Water.

#### RT primers for library preparation

Dissolve RT primers to make 10 μM stock solutions in Nuclease-Free Water. To enhance sequence diversity during sequencing, prepare 4 µM solutions comprising pairs of RT primers (2 µM each) that either exhibit staggered binding to the reporter or incorporate a varying number of inserted untemplated nucleotides (Fig. 2E, **Supporting Information:** Fig. S1 for all sequences). RT primer stock and working solutions can be stored at -20°C for a minimum of one year.

#### 10X Salt Buffer

Prepare a solution containing 150 mM Tris pH 7.5, 1 M NaCl (Thermo Fisher Scientific, Cat. No. AM9760G), and 100 mM MgCl_2_ (Thermo Fisher Scientific, Cat. No. AM9530G), adjusting to the final volume with Nuclease-Free Water.

#### Spike-in mastermix

Clone and synthesize each spike-in following the **Support Protocol 1**. Pool spike-ins to desired concentrations. Our pool consisted of 1, 5, 25, 50, and 125 fg/µL for each spike-in 1 to 5, respectively, in a final volume of 500 µL with Nuclease-Free Water (see Table 2 for an example of dilution scheme). Bring volume up to 1 mL with an additional 500 µL of Nuclease-Free Water, then freeze this mastermix in aliquots at -80°C and use at 2 µL/mL of TRIzol at the extraction step (*e*.*g*., prepare 15 mL of TRIzol + 30 µL of spike-in mastermix for 15 samples, and use 1 mL per sample).

**Table 2.**
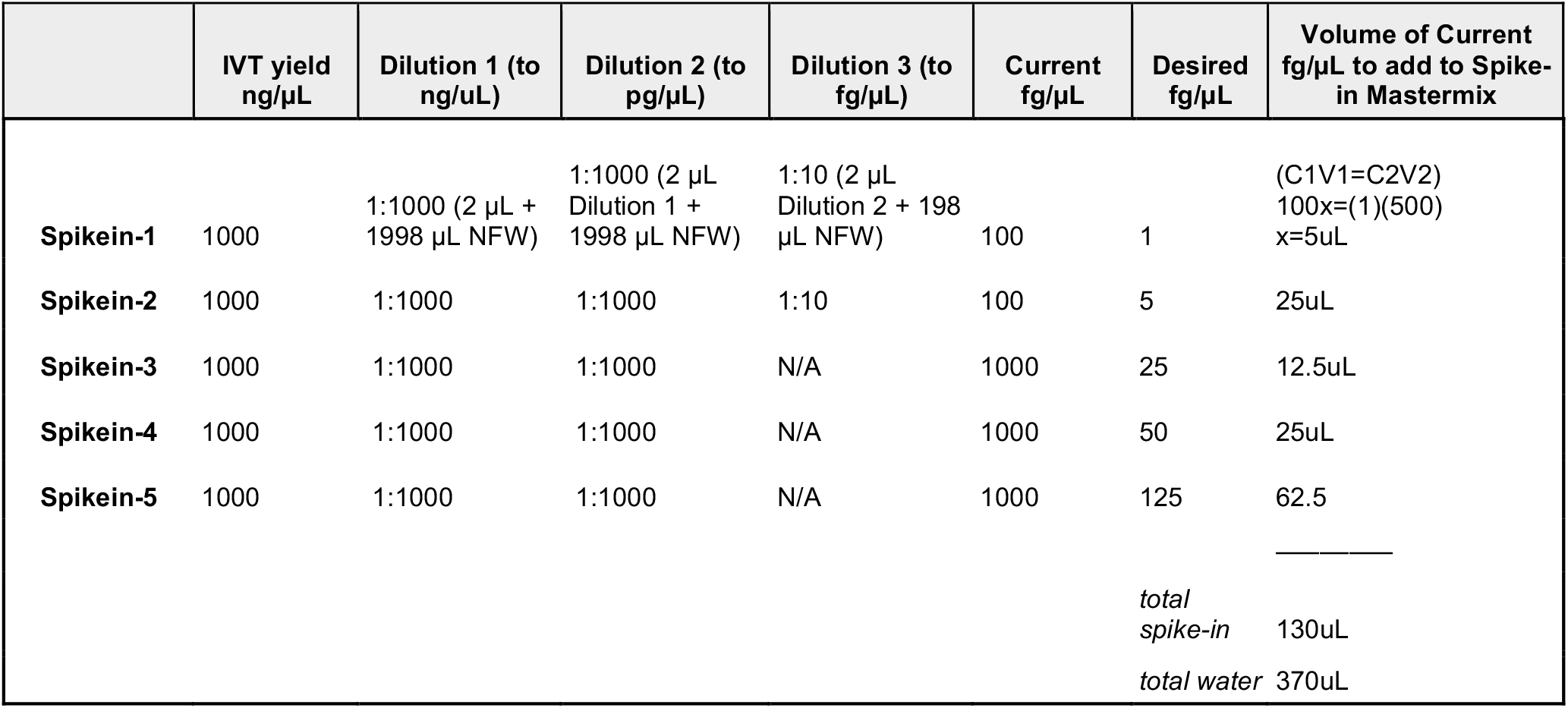
Example dilution math for spike-in mastermix.

#### 10× TAE buffer

Dissolve 48.5 g of Tris in 800 mL distilled water. Add 11.4 mL glacial acetic acid and 20 mL 0.5 M EDTA (pH 8.0) and complete to 1 L using distilled water for a final concentration of 400 mM Tris, 200 mM acetic acid and 10 mM EDTA. Store long term at room temperature.

### COMMENTARY

#### Background Information

The NaP-TRAP procedure results in the measurement of reporter abundance in both the input and pulldown fractions during an immunocapture experiment. The input sample values can be directly examined to observe changes in reporter abundance over time or under different conditions. Due to the design of NaP-TRAP, the ratio of the pulldown to input fractions indicates the translation of the ORF that encodes the epitope targeted for pulldown (Fig. 1B). NaP-TRAP can be conducted with either a small number of reporters or complex reporter libraries, utilizing qPCR or high-throughput sequencing as the respective readouts (Fig. 1A). Here, we outlined typical results from experiments conducted in zebrafish embryos and HEK293T mammalian cells for both throughput methods. A detailed workflow of the bioinformatic analysis is accessible on our GitHub (Madhavan and Strayer, 2025) (Fig. 4).

#### Critical Parameters

This protocol is designed to assess how mRNA regulatory elements influence translation in zebrafish embryos and cultured cell lines. While our initial study focused on the 5′-UTR regulatory landscape, NaP-TRAP can be adapted to interrogate any mRNA region, assuming proper experimental design. We have used NaP-TRAP to examine elements across the full mRNA, from the cap structure and uORFs in the 5′-UTR, to codon optimality in the coding sequence (CDS), microRNA binding sites in the 3′-UTR, and poly(A) tail length, in zebrafish embryos. The results underscore NaP-TRAP’s extensive adaptability in investigating various mechanisms of translational regulation. Here, we outline essential factors to consider for the effective implementation of NaP-TRAP in various experimental contexts.

#### Reporter design: Basics

Each NaP-TRAP reporter must contain two essential elements: an N-terminal epitope tag and a C-terminal protein degron domain within the open reading frame (ORF) used for translation readout. We recommend using a 3xFLAG epitope tag when possible, though we have successfully used MYC and HA tags. For the degron, we advise using the PEST domain (Rechsteiner and Rogers, 1996), as it effectively reduces protein half-life in most tested cell lines and does not require additional components for activity. If the PEST domain is not suitable for your system, the auxin-inducible degron (AID) system offers an alternative; however, this approach requires co-expression of the TIR1 protein and auxin treatment.

All NaP-TRAP experiments to date have employed *in vitro* transcribed mRNA reporters generated from PCR products or digested plasmids using T7 or SP6 RNA polymerase. For 3′-end processing, we have tested two approaches: (1) cytoplasmic cleavage and polyadenylation using the SV40 polyadenylation signal, and (2) hard-encoded poly(A) tails introduced via a reverse PCR primer. It is important to note that the SV40 signal functions only in zebrafish embryos (Collas and Aleström, 1997) and results in lower translation levels compared to hard-encoded poly(A) tails (Strayer et al., 2024). Importantly, the SV40 approach does not work in human cells when RNA is directly transfected. Therefore, we recommend generating NaP-TRAP reporters by PCR amplification, adding the promoter (*e*.*g*., SP6 or T7) in the forward primer and a defined poly(A) tail (stretch of Ts) in the reverse primer. The PCR template can be a plasmid for individual reporters or assembled via PCR for constructing complex reporter libraries (Fig. 2A-B). This strategy produces reporters with precisely defined sequences from the 5′ to 3′ ends, effectively avoiding artifacts that may arise during transcription or pre-mRNA processing.

The next consideration for reporter design involves the number of reporters to be tested simultaneously and the method for quantifying enrichment in the input and pulldown samples. For small numbers of reporters, we recommend quantitative PCR (qPCR). For larger libraries, Illumina sequencing provides a scalable, high-throughput readout.

#### Reporter design: qPCR readout

A qPCR readout is well-suited for experiments involving a small pool of reporters, typically up to a dozen, depending on the throughput capacity of the qPCR platform. Following RNA extraction from both input and immunoprecipitated samples, reporters are reverse transcribed using random hexamers. The resulting cDNA is diluted and used as the template in qPCR reactions with primer pairs specific to each reporter. For optimal specificity and simplicity, one primer should anneal to a constant region shared by all reporters, while the other should target a unique sequence within the variable region (Fig. 2C). This strategy allows multiple reporters to be assessed in the same NaP-TRAP experiment, reducing variability and enabling direct comparison of translational activity across reporters within a single immunoprecipitation reaction.

#### Reporter design: Illumina sequencing readout

Illumina sequencing is the preferred readout for NaP-TRAP when conducting MPRAs, enabling the analysis of thousands of sequences in a single experiment. Advances in oligo synthesis now allow routine, cost-effective production of high-quality pools of varying lengths and sizes from commercial vendors. Each oligo in the pool contains a variable region, or insert, flanked by two fixed sequences common to all oligos. These fixed sequences are essential for PCR amplification of the oligo pool, which is then assembled via PCR into the full NaP-TRAP reporter construct to serve as a template for *in vitro* transcription (Fig. 2D). The insert length and total number of oligos can be tailored to match the experimental design and the regulatory sequence space being interrogated. For example, we initially applied NaP-TRAP to characterize the 5′-UTR regulatory landscape during zebrafish embryogenesis. We designed 170-nucleotide oligos comprising a 124-nucleotide variable insert flanked by 24- and 22-nucleotide fixed sequences at the 5′ and 3′ ends, respectively. Both fixed sequences should be distinct from each other to ensure correct polarity in the final assembly reaction. The inserts were derived from 1,775 endogenous 5′-UTRs and six viral IRES elements, sampled using a sliding window of 124 nucleotides with a step size of 25 nucleotides (Fig. 2D), resulting in a library of 11,088 oligos. It is important to include internal controls in each library that contain well-characterized regulatory elements relevant to the system being studied (see **NaP-TRAP controls** section below). In our 5′-UTR library, we have multiple inserts with uAUGs and oORFs, which are known global repressors of translation, to serve as benchmarks.

When designing a NaP-TRAP library, it is essential to clearly define which mRNA regions will vary and which will remain constant. In our 5′-UTR screen, we kept the 5′-UTR length and the Kozak sequence constant to specifically identify novel regulatory elements within the variable regions, as both are strong determinants of translation. If our primary goal had been to predict translation initiation as modulated by full-length endogenous 5′-UTRs, a more appropriate approach would have been to include entire 5′-UTRs constrained to fit within the oligo length limits of the synthesis pool. In that case, both 5′-UTR length and Kozak sequence would vary across reporters to reflect their native sequence contexts. This strategy was used by (Reimão-Pinto et al., 2023) to investigate how 5′-UTR sequences influence translation in zebrafish embryos.

#### Pulldown and spike-ins

Spike-ins can be introduced at various stages of the NaP-TRAP workflow to serve as internal normalizers and reduce technical variability. For RT-qPCR-based readouts, we recommend co-delivering a non-FLAG-tagged mRNA, such as dsRed, alongside NaP-TRAP reporters during transfection or micro-injection. qPCR measurements of this spike-in can then be used to normalize both input and pulldown samples. For Illumina sequencing readouts, we suggest adding spike-ins directly into the TRIzol reagent used for RNA extraction from input and pulldown samples. These spike-ins should mirror the structure of the NaP-TRAP library, including fixed regions and insert length, but only containing insert sequences absent from the experimental library to ensure unambiguous identification. To enhance sequence complexity and manage noise across various sequencing depths, we recommend using a mix of five distinct *in vitro* transcribed spike-ins that span a broad range of concentrations. The optimal amount of spike-in RNA depends on the sample type (*e*.*g*., zebrafish embryos vs. cultured cells) and total RNA yield. As a general guideline, spike-ins should account for 0.1–2% of total sequencing reads.

#### Considerations for Illumina library sequencing preparation

Below are key considerations for generating libraries compatible with Illumina’s dual-indexed paired-end sequencing workflow. Sequencing quality is highly dependent on sequence diversity, particularly in the initial cycles of read 1. To maximize heterogeneity in read 1, we recommend incorporating a region recognized by the read 1 sequencing primer (i5) directly upstream of the variable insert, ensuring that read 1 begins precisely at the first base of the insert (Fig. 2E). A region recognized by the read 2 sequencing primer (i7) sequence should be placed at the 5′ end of the reverse transcription (RT) primer, which binds downstream of the insert (Fig. 2E). The full RT primer design, from 5′ to 3′, includes: the i7 region, a 4-nucleotide anchor, a 10-nucleotide unique molecular identifier (UMI), another 4-nucleotide anchor, a 6-nucleotide custom barcode, and a reverse complement sequence recognizing the reporter (Fig. 2E). This design allows the resulting cDNA to be directly amplified using Illumina TruSeq dual-indexing primer P5 and P7 (Fig. 2E). To reduce PCR samples and variability across replicates, we recommend assigning a distinct custom barcode to each biological replicate and pooling them immediately after RT for a shared PCR amplification (Fig. 2F).

In situations where it is not possible to place the i5 sequence directly upstream of the insert, an alternative approach involves using an extended forward PCR primer containing the complete P5 primer sequence followed by a region that anneals near the insert (**Supporting Information**: Fig. S1A). To improve sequence diversity in read 1 under this setup, use a mix of three versions of the P5 primer, each with 0, 1, or 2 untemplated nucleotides inserted between the i5 and annealing regions. This strategy creates sequencing reads in staggered frames, increasing nucleotide diversity during the early cycles (Fig. 2E and **Supporting Information**: Fig. S1B). It is essential that read 1 spans enough of the variable region to uniquely map each read to its corresponding insert. A similar staggering approach can be applied to read 2 by inserting untemplated nucleotides between the custom barcode and the annealing sequence in the RT primer (Fig. 2E and **Supporting Information**: Fig. S1B). For optimal sequencing performance, especially when using fixed upstream sequences, adjust the ratio of PhiX to NaP-TRAP libraries during sequencing. Using the staggered-frame strategy described above, we obtained high-quality reads on a MiSeq platform with as little as 40% PhiX. Similarly, co-sequencing NaP-TRAP libraries with other complex libraries on platforms like NovaSeq improves sequence diversity and read quality.

#### Considerations for sequencing depth

The depth of sequencing is influenced by the complexity of the reporter library and the number of amplification cycles, with a higher number of cycles leading to more PCR duplicates which will be eliminated before analysis. For NaP-TRAP libraries comprising several dozen reporters, sufficient coverage was achieved using in-house MiSeq sequencing, as detailed in the step-by-step procedure included in this protocol. However, for more complex libraries, such as the zebrafish 5’-UTR library containing 11,088 reporters, sequencing on higher throughput platforms, such as NovaSeq or NextSeq 2000, is necessary to ensure adequate coverage. It is advisable to aim for a minimum of 100 reads per insert in each replicate (*e*.*g*., input replicate 1, pulldown replicate 1, etc.). We recommend exceeding this number by at least a factor of 10 to incorporate as many reporters as possible, particularly those with lower abundance, in the analysis. If higher throughput platforms will be used, we suggest conducting a MiSeq run regardless to check the quality of read distribution across reporters and to determine the percentage of reads mapping to spike-ins for each sample.

#### NaP-TRAP controls

When implementing NaP-TRAP for the first time as a new user or in a new system (*e*.*g*., a different mammalian cell line), we recommend validating the protocol using reporters that contain well-characterized regulatory elements. In our hands, control reporters incorporating features such as mRNA cap-analogs, upstream open reading frames (uORFs), codon optimality motifs, microRNA binding sites, morpholino binding sites (Strayer et al., 2024), and varying poly(A) tail lengths have been effective for this purpose (Fig. 1E). A particularly robust and versatile validation strategy involves comparing NaP-TRAP reporters with or without an uORF that overlaps with the main ORF (Fig. 1E). Given the strong and global repressive effects of overlapping ORFs on translation, this comparison provides a reliable functional benchmark in most systems (**Supporting Information**: Fig. S2). These validations can be performed cost-effectively using qPCR as the readout (see **Supporting Information**: Table S2 for example qPCR analysis). When designing NaP-TRAP libraries for sequencing, it is important to include control sequences that have already been validated using the NaP-TRAP qPCR workflow. These controls should be located within the same mRNA region as your experimental inserts and be compatible with the library preparation strategy (Fig. 2E). It is essential to ensure that the selected regulatory elements are functionally active and well-suited to the specific biological system being studied.

#### Plasmid validation

To test the impact of 5’-UTR elements on translation regulation, an overlapping ORF was cloned into the 5’-UTR of the 3xFLAG-GFP open reading frame. To examine the effects of optimal versus non-optimal codons, constructs were adapted from (Bazzini et al., 2016), in which insertion of a single nucleotide switches an ORF from non-optimal to optimal codon composition. A 3xFLAG tag was inserted after the ORF initiation codon and upstream of the nucleotide insertion site that alters codon optimality. To assess 3’-UTR effects on translation, wild-type or mutated 3x miRNA-430 sites (Bazzini et al., 2012) were cloned downstream of the 3xFLAG-GFP open reading frame. To evaluate the role of poly(A)-tail length, the 3xFLAG-GFP construct lacking an upstream AUG was PCR-amplified using reverse primers encoding poly(A) tails of defined lengths, and the resulting PCR products were used for in vitro transcription to generate reporters with variable poly(A) tails.

#### Troubleshooting Table

Table 3 discusses common problems with the protocol, their causes, and potential solutions.

**Table 3.**
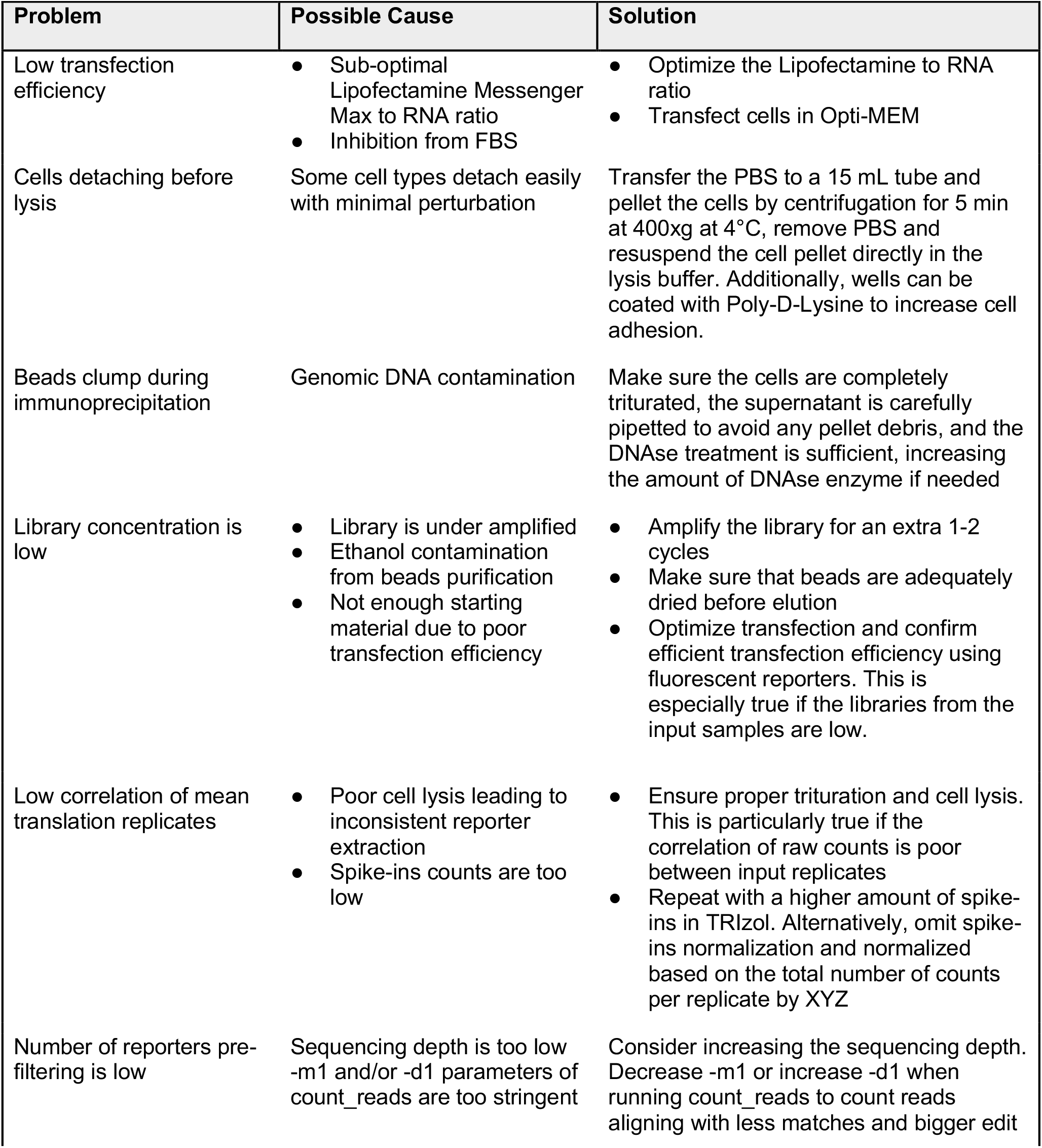
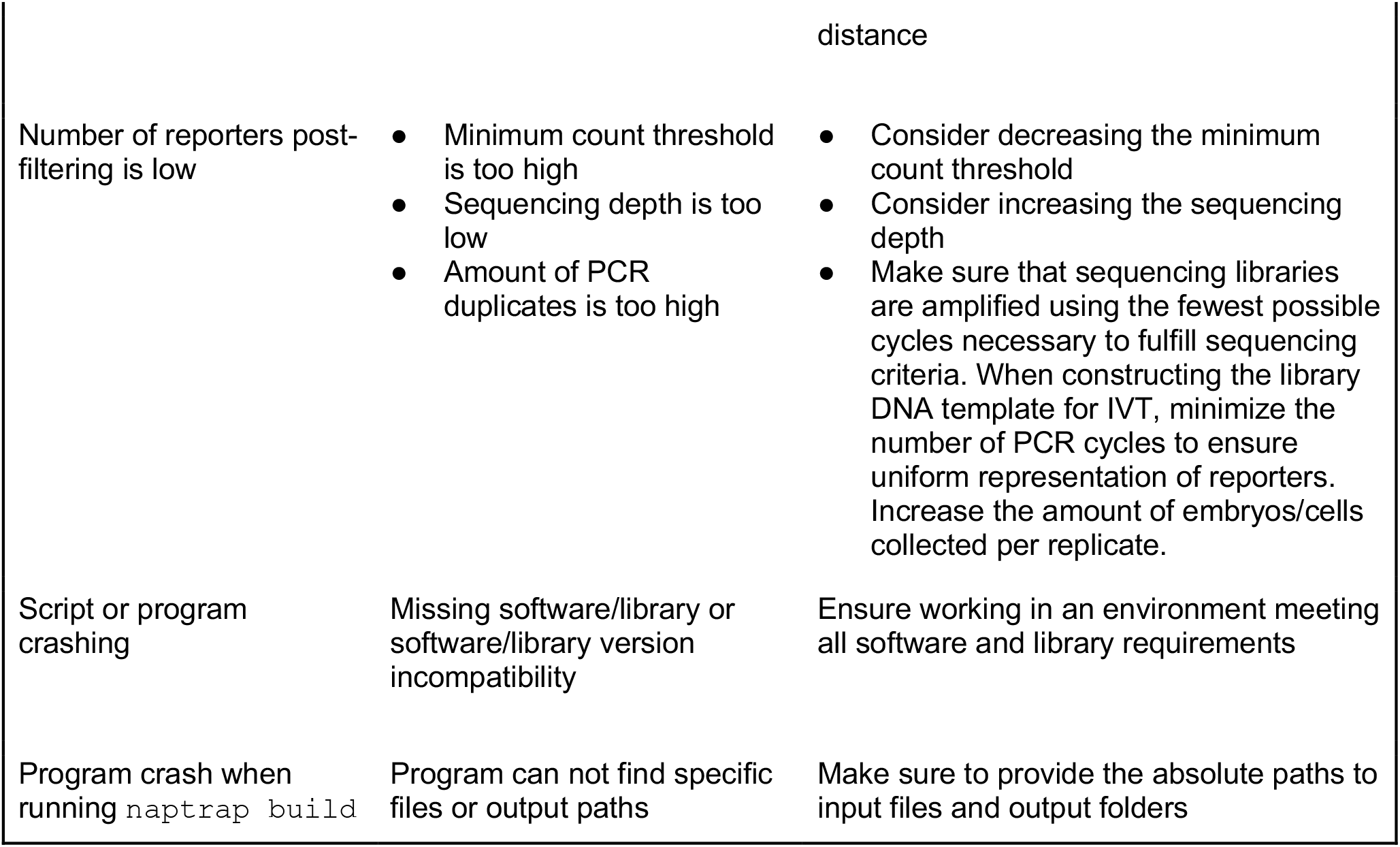
Troubleshooting Guide for NaP-TRAP.

#### Understanding Results

For NaP-TRAP experiments using qPCR as a readout, we included examples of qPCR analysis sample sheets for experiments involving reporters with or without an oORF, as well as those with or without miR-430 sites in their 3’-UTR (see **Supporting Information**: Table S2). Translation values are calculated using the 2^-(ΔΔCq)^ method as detailed above in **Alternate Protocol 2**. For the with and without oORF experiment, reporters were transfected (Fig. 1D) or microinjected (Fig. 1E) separately because they differ by only a few nucleotides, which is insufficient for specific amplification during qPCR if co-injected. Inversely, reporters with or without miR-430 (Fig. 1A and 1E) were microinjected together, as their sequences differ enough to allow specific amplification with designated qPCR primers. In both experimental setups, NaP-TRAP translation values reflect the overall repressive impact of oORFs across various systems (Fig. 1D-E) and the temporal repression of miR-430 in zebrafish embryos (Fig. 1E). Importantly, we demonstrated the adaptability of NaP-TRAP in assessing the regulatory activity of different elements throughout the mRNA molecule. This includes the activating influence of the mRNA cap-structure, the positive and negative effects of optimal and non-optimal codons, respectively, and the enhancement of translation by a longer poly(A) tail in zebrafish embryos (Fig. 1E).

For NaP-TRAP experiments using high throughput sequencing, we provided examples of MPRA analyses in both zebrafish embryos and HEK293T mammalian cells (Fig. 1C-E) (Strayer et al., 2024). The number of reporters passing the minimum read coverage and included in the analysis depends on: i) the uniform representation of each reporter in the input samples (Fig. 3A), ii) the depth of sequencing reads, and iii) the number of cycles used for sequencing library amplification, which affects the removal of PCR duplicates. For our library containing 10,088 reporters, with sequencing read depths ranging from X-Y per sample, libraries amplified between 12-18 PCR cycles, and a minimum requirement of 100 reads per insert in all input replicates, we obtained between X to Y reporters passing our filter. Reducing the minimum read requirement is possible, but it involves a tradeoff between the number of reporters passing the filter and the signal noise that impacts the correlation between replicates. With our chosen parameters, we achieved high correlations (from 0.91 to 0.97) in translation values across replicates (Fig. 3B-C).

Additionally, the provided scripts can generate a histogram of translation values for a single condition and conduct a kmer enrichment analysis for reporters exhibiting low (orange) and high (blue) translation (Fig. 3D-E). Here, there is a clear enrichment for AUG-containing kmers and U-rich kmers in reporters exhibiting low and high translation, respectively, in zebrafish embryos at 2 hpf (Fig. 3D-E). When comparing translation between two conditions, such as 2 and 6 hpf in zebrafish, scatter plots can be produced, categorizing reporters into four groups: i) low (orange) or ii) high (blue) translation at both stages, iii) low at 2 hpf and high at 6 hpf (green), and iv) high at 2 hpf and low at 6 hpf (pink) (Fig. 3F). To gain deeper insights into the elements influencing translation between these stages, a kmer enrichment analysis can be conducted for each group (Fig. 3G). For all enrichment analyses, besides the enrichment plots (Fig. 3C), the pipeline also provides tables with the corresponding raw values.

Finally, our analytical pipeline outputs various tables (Fig. 4) that include the insert sequence, raw count, normalized count, and translation value for each reporter, both before and after filtering. These tables offer flexibility and serve as excellent starting points for further downstream analyses.

#### Time Considerations

The full NaP-TRAP workflow is visualized in Fig. 1A. Library Assembly and IVT timing (**Basic Protocol 1**) timing depends on whether reporter constructs will be cloned (**Support Protocol 1**) before being amplified for IVT. Each PCR reaction and gel extraction can be completed in ∼2 hrs of hands-on time and 4 hrs total. Bacterial cloning adds an additional 1 hr of hands-on time, 2 overnight incubations for transformation and single colony isolation, as well as wait-time for Sanger-sequencing validation. Once reporters are confirmed, IVT takes ∼4 hrs with ∼30 min hands-on time, including quality check by agarose gel.

mRNA reporter delivery takes 2-3 days for set-up depending on the model system. **Basic Protocol 2** begins one day prior to planned injections to set crossing tanks of zebrafish embryos (∼1 hr hands-on). Crossing takes ∼1 hr of hands-on time to change water and collect embryos, and injection should be completed within 35 min to avoid passing the window of 1-cell stage. Embryos can be collected at any desired timepoint, flash-freezing and stored long-term at -80°C. **Alternate Protocol 1** delivery into cells requires several days in advance to thaw cells, and passage for optimal density at transfection (∼1 hr hands-on time). On the day of transfection, it takes ∼10 min to prepare reagents and transfect cells, then 1 hr of incubation. Next, it takes ∼30 min of hands-on time to collect cells post-transfection, which can be either frozen or kept on ice for next steps.

NaP-TRAP pulldown takes ∼3 hrs (∼30 min hands-on) and is followed by RNA isolation which takes ∼2 hrs plus 1 hrs wait-time (**Basic Protocol 3**). Isolated RNA from pulldown and input are then reverse transcribed, and synthesized cDNA is amplified for high-throughput sequencing library preparation (**Basic Protocol 4**; ∼4 hrs total, ∼1 hr hands-on), plus an additional 2 days of wait time for MiSeq sequencing (∼30 min hands-on). Alternatively, RNA can be reverse transcribed and analyzed by RT-qPCR (**Alternate Protocol 2**; ∼4 hr total, ∼1 hr hands-on, including data analysis). The time required for computational analysis of sequencing datasets depends on experimental parameters, sequencing depth, available computational resources, and user experience, but typically involves ∼6-7 hrs of runtime with ∼1-2 hrs of user input for MiSeq runs analysis and ∼2-3 days of runtime with 2-3 hrs of user input for deeper NovaSeq runs (**Basic Protocol 5**).

## ACKNOWLEDGMENTS

We thank Srikar Krishna from the Giraldez lab at Yale for feedback on the manuscript, and all members of the Beaudoin lab for feedback and support. We would like to acknowledge our funding sources: National Institutes of Health grant R35 GM146883 (J.-D.B) and NHGRI T32 HG010463 (A.Z.S).

## DATA AVAILABILITY STATEMENT

The data used to generate the anticipated results in the MPRA experiments were originally published in Strayer *et al*. All sequencing data are available at NCBI under the BioProject ID PRJNA1188270 (see **Supporting Information**: Table S3 for details). Additional specific details can be obtained from the corresponding author upon reasonable request.

## SUPPORTING INFORMATION

| Filename | Description |
| --- | --- |
| TableS1_oligos_plasmids.xlsx | *Table S1: Excel table containing all relevant sequences for oligos and plasmids.* |
| TableS2_qPCR_example_sheets.xlsx | *Table S2: Excel table with example output and calculations for NaP-TRAP-qPCR.* |
| TableS3_Sequencing_samples.xlsx | *Table S3: Excel table containing all sequencing sample IDs and descriptions relevant to this protocol.* |
| SupportingFigures.pdf | *Figure S1. Additional considerations for NaP-TRAP sequencing library designs.*<br>*Figure S2. Orthogonal validation of reporter repression by an oORF in mammalian cells.* |

Original paper first reporting NaP-TRAP and its use to define the regulatory grammar of 5′ UTR-mediated translation in zebrafish embryos and mammalian systems.

## INTERNET RESOURCES

Code used for the NaP-TRAP analysis is accessible via the Beaudoin lab GitHub (https://github.com/beaudoinlab/nap_trap_protocol), and is available for reuse.

## LITERATURE CITED

Akirtava, C., and McManus, C. J. 2021. Control of translation by eukaryotic mRNA transcript leaders - insights from high-throughput assays and computational modeling. Wiley interdisciplinary reviews. RNA 12:e1623.

Bazzini, A. A., Del Viso, F., Moreno-Mateos, M. A., Johnstone, T. G., Vejnar, C. E., Qin, Y., Yao, J., Khokha, M. K., and Giraldez, A. J. 2016. Codon identity regulates mRNA stability and translation efficiency during the maternal-to-zygotic transition. The EMBO journal 35:2087–2103.

Bazzini, A. A., Lee, M. T., and Giraldez, A. J. 2012. Ribosome Profiling Shows That miR-430 Reduces Translation Before Causing mRNA Decay in Zebrafish. Science 336:233–237.

Boswell, C. W., Hoppe, C., Sherrard, A., Miao, L., Kojima, M. L., Martino, P., Zhao, N., Stasevich, T. J., Nicoli, S., and Giraldez, A. J. 2025. Genetically encoded affinity reagents are a toolkit for visualizing and manipulating endogenous protein function in vivo. Nature Communications 16:5503.

Byeon, G. W., Cenik, E. S., Jiang, L., Tang, H., Das, R., and Barna, M. 2021. Functional and structural basis of extreme conservation in vertebrate 5′ untranslated regions. Nature Genetics 53:729–741.

Calvo, S. E., Pagliarini, D. J., and Mootha, V. K. 2009. Upstream open reading frames cause widespread reduction of protein expression and are polymorphic among humans. Proceedings of the National Academy of Sciences of the United States of America 106:7507–7512.

Chassé, H., Boulben, S., Costache, V., Cormier, P., and Morales, J. 2017. Analysis of translation using polysome profiling. Nucleic Acids Research 45:e15.

Cleary, J. D., and Ranum, L. P. W. 2013. Repeat-associated non-ATG (RAN) translation in neurological disease. Human Molecular Genetics 22:R45–51.

Collas, P., and Aleström, P. 1997. Rapid targeting of plasmid DNA to zebrafish embryo nuclei by the nuclear localization signal of SV40 T antigen. Molecular Marine Biology and Biotechnology 6:48–58.

Cuperus, J. T., Groves, B., Kuchina, A., Rosenberg, A. B., Jojic, N., Fields, S., and Seelig, G. 2017. Deep learning of the regulatory grammar of yeast 5′ untranslated regions from 500,000 random sequences. Genome Research 27:2015–2024.

Dao, K., Jungers, C. F., Djuranovic, S., and Mustoe, A. M. 2025. U-rich elements drive pervasive cryptic splicing in 3’ UTR massively parallel reporter assays. Nature Communications 16:6844.

Diagrams created in BioRender. Smith, J. (2025). BioRender.com/c248457.

Fabbri, L., Chakraborty, A., Robert, C., and Vagner, S. 2021. The plasticity of mRNA translation during cancer progression and therapy resistance. Nature Reviews. Cancer 21:558–577.

Fagre, C., and Gilbert, W. 2024. Beyond reader proteins: RNA binding proteins and RNA modifications in conversation to regulate gene expression. Wiley interdisciplinary reviews. RNA 15:e1834.

Gilliot, P.-A., and Gorochowski, T. E. 2023. Effective design and inference for cell sorting and sequencing based massively parallel reporter assays. Bioinformatics (Oxford, England) 39:btad277.

Gingold, H., and Pilpel, Y. 2011. Determinants of translation efficiency and accuracy. Molecular Systems Biology 7:481.

Gupta, A., and Beaudoin, J.-D. 2021. Extreme conservation encodes the structural dynamics and function of 5’ UTRs. Nature Genetics 53:591–592.

Heiman, M., Schaefer, A., Gong, S., Peterson, J. D., Day, M., Ramsey, K. E., Suárez-Fariñas, M., Schwarz, C., Stephan, D. A., Surmeier, D. J., et al. 2008. A translational profiling approach for the molecular characterization of CNS cell types. Cell 135:738–748.

Hentze, M. W., Castello, A., Schwarzl, T., and Preiss, T. 2018. A brave new world of RNA-binding proteins. Nature Reviews. Molecular Cell Biology 19:327–341.

Ingolia, N. T., Ghaemmaghami, S., Newman, J. R. S., and Weissman, J. S. 2009. Genome-wide analysis in vivo of translation with nucleotide resolution using ribosome profiling. Science (New York, N.Y.) 324:218–223.

Ingolia, N. T., Hussmann, J. A., and Weissman, J. S. 2019. Ribosome Profiling: Global Views of Translation. Cold Spring Harbor Perspectives in Biology 11:a032698.

Johnstone, T. G., Bazzini, A. A., and Giraldez, A. J. 2016. Upstream ORFs are prevalent translational repressors in vertebrates. The EMBO journal 35:706–723.

Kelleher, R. J., and Bear, M. F. 2008. The autistic neuron: troubled translation? Cell 135:401–406.

Kingston, E. R., Blodgett, L. W., and Bartel, D. P. 2022. Endogenous transcripts direct microRNA degradation in Drosophila, and this targeted degradation is required for proper embryonic development. Molecular Cell 82:3872–3884.e9.

Langmead, B., and Salzberg, S. L. 2012. Fast gapped-read alignment with Bowtie 2. Nature Methods 9:357–359.

Madhavan, S., and Strayer, E. 2025. beaudoinlab/nap_trap_protocol. Available at: https://github.com/beaudoinlab/nap_trap_protocol.

Medina-Muñoz, S. G., Kushawah, G., Castellano, L. A., Diez, M., DeVore, M. L., Salazar, M. J. B., and Bazzini, A. A. 2021. Crosstalk between codon optimality and cis-regulatory elements dictates mRNA stability. Genome Biology 22:14.

Meijer, H. A., and Thomas, A. A. M. 2002. Control of eukaryotic protein synthesis by upstream open reading frames in the 5’-untranslated region of an mRNA. The Biochemical Journal 367:1–11.

Mikl, M., Eletto, D., Nijim, M., Lee, M., Lafzi, A., Mhamedi, F., David, O., Sain, S. B., Handler, K., and Moor, A. E. 2022. A massively parallel reporter assay reveals focused and broadly encoded RNA localization signals in neurons. Nucleic Acids Research 50:10643–10664.

Niederer, R. O., Rojas-Duran, M. F., Zinshteyn, B., and Gilbert, W. V. 2022. Direct analysis of ribosome targeting illuminates thousand-fold regulation of translation initiation. Cell Systems 13:256–264.e3.

Rabani, M., Pieper, L., Chew, G.-L., and Schier, A. F. 2017. A Massively Parallel Reporter Assay of 3′ UTR Sequences Identifies In Vivo Rules for mRNA Degradation. Molecular Cell 68:1083–1094.e5.

Rechsteiner, M., and Rogers, S. W. 1996. PEST sequences and regulation by proteolysis. Trends in Biochemical Sciences 21:267–271.

Reimão-Pinto, M. M., Castillo-Hair, S. M., Seelig, G., and Schier, A. F. 2023. The regulatory landscape of 5’ UTRs in translational control during zebrafish embryogenesis. bioRxiv: The Preprint Server for Biology:2023.11.23.568470.

Schmidt, T., Turnbull, A. P., Gupta, A., Waldron, J. A., Pollock, K., Munro, J., Nepravishta, R., Herviou, P., Dąbrowska, A., Hodge, K., et al. 2025. The differential mechanisms of eIF4A1-mediated translational activation instructed by distinct RNA features. 2025.03.24.644895. Available at: https://www.biorxiv.org/content/10.1101/2025.03.24.644895v1 [Accessed November 11, 2025].

Shi, X., Sun, M., Wu, Y., Yao, Y., Liu, H., Wu, G., Yuan, D., and Song, Y. 2015. Post-transcriptional regulation of long noncoding RNAs in cancer. Tumour Biology: The Journal of the International Society for Oncodevelopmental Biology and Medicine 36:503–513.

Siegel, D. A., Le Tonqueze, O., Biton, A., Zaitlen, N., and Erle, D. J. 2022. Massively parallel analysis of human 3’ UTRs reveals that AU-rich element length and registration predict mRNA destabilization. G3 (Bethesda, Md.) 12:jkab404.

Strayer, E. C., Krishna, S., Lee, H., Vejnar, C., Neuenkirchen, N., Gupta, A., Beaudoin, J.-D., and Giraldez, A. J. 2024. NaP-TRAP reveals the regulatory grammar in 5’UTR-mediated translation regulation during zebrafish development. Nature Communications 15:10898.

Vejnar, C. E., and Giraldez, A. J. 2020. LabxDB: versatile databases for genomic sequencing and lab management. Bioinformatics (Oxford, England) 36:4530–4531.

Xiang, K., Ly, J., and Bartel, D. P. 2024. Control of poly(A)-tail length and translation in vertebrate oocytes and early embryos. Developmental Cell 59:1058–1074.e11.

Zhao, W., Pollack, J. L., Blagev, D. P., Zaitlen, N., McManus, M. T., and Erle, D. J. 2014. Massively parallel functional annotation of 3’ untranslated regions. Nature Biotechnology 32:387–391.

Zuccotti, P., and Modelska, A. 2016. Studying the Translatome with Polysome Profiling. Methods in Molecular Biology (Clifton, N.J.) 1358:59–69.

