## Supplementary Figures for "NaP-TRAP: A versatile and accessible workflow to dissect principles of translational regulation and mRNA stability"

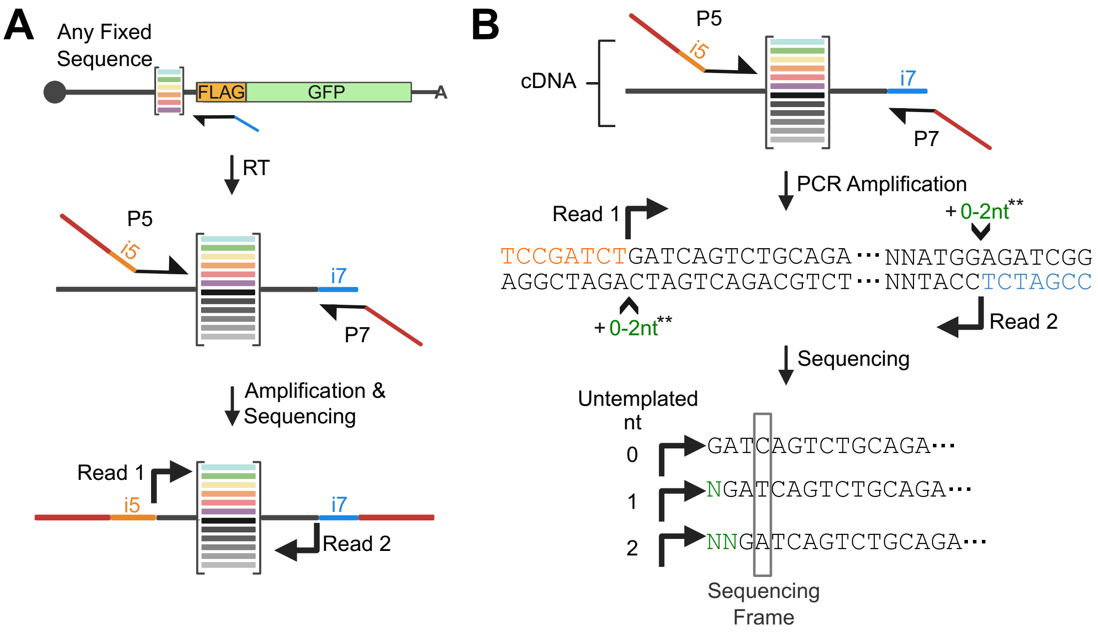


**Supporting Figure S1.** Additional considerations for NaP-TRAP sequencing library designs. (**A**) If it is not feasible to encode part of the i5 sequence (orange) immediately upstream of the insert, use a modified PCR primer in which a sequence complementary to the fixed region upstream of the insert (black) is appended to the 3' end of the P5-i5 adapter sequence. (**B**) In this configuration, to increase sequence diversity at the start of read 1, we recommend using a mixture of modified primers that contain 0-2 additional nucleotides (green) inserted between the i5 (orange) and binding sequence (black). This staggered-primer strategy increases heterogeneity within the upstream fixed sequence during sequencing. A similar staggered-primer strategy can also be applied for the RT primer to increase sequence diversity at the start of read 2 (see also Fig. 2E).

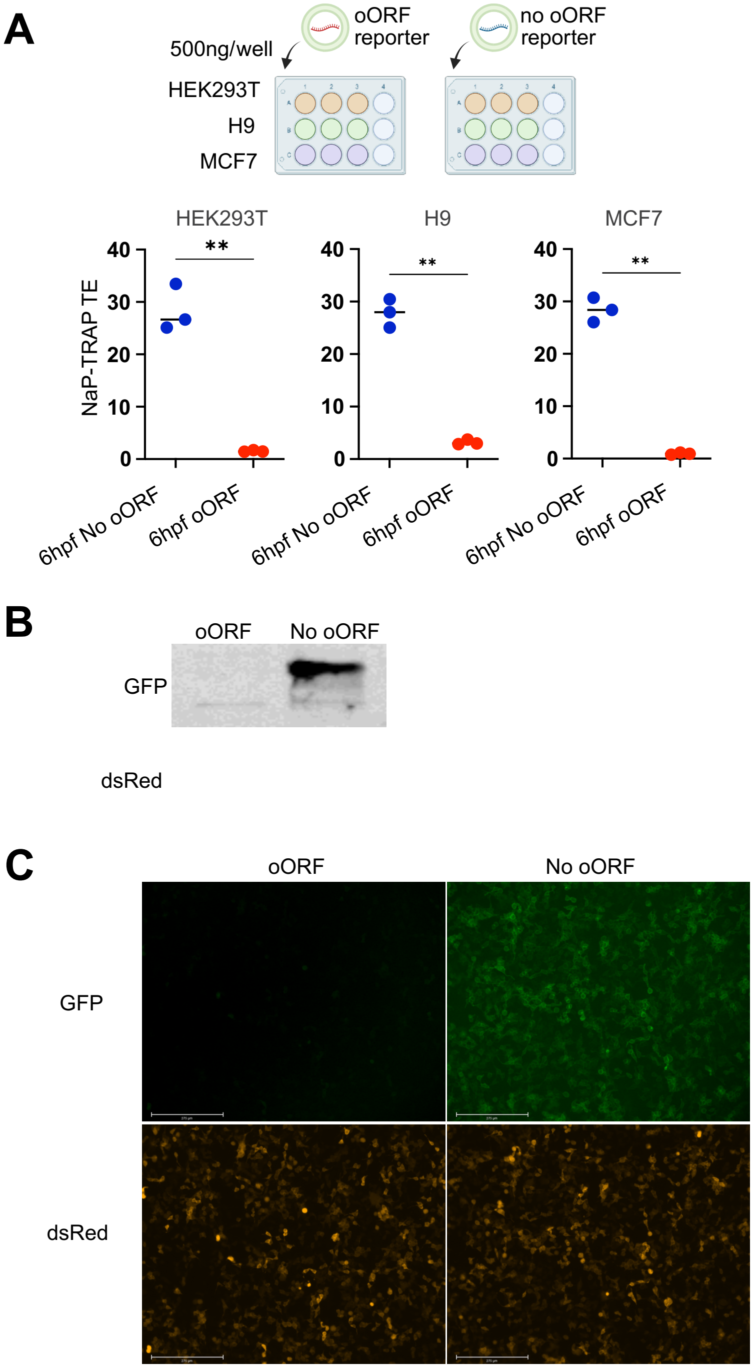


**Supporting Figure S2.** Orthogonal validation of reporter repression by an oORF in mammalian cells. (**A**) NaP-TRAP translation efficiency (TE) of reporters with or without an overlapping ORF in HEK293T cells. (**B**) GFP and dsRed (control) protein expression measured by Western blot for the same reporters in HEK293T cells. (**C**) Fluorescence microscopy of NaP-TRAP reporters with and without overlapping ORF (GFP), with dsRed co-delivered as a transfection control in HEK293T cells. All three approaches capture the repressive effect of the overlapping ORF on translation.
